# SWANS: A highly configurable analysis pipeline for single-cell and single-nucleus RNA-sequencing data

**DOI:** 10.1101/2025.05.14.654073

**Authors:** Katherine Beigel, Eric Wafula, Dana V. Mitchell, Steven J. Pastor, Michelle Gong, Robert O. Heuckeroth, Julio C. Ricarte-Filho, Aime T. Franco, Erin R. Reichenberger

## Abstract

**Background:** Single-cell RNA sequencing (scRNA-seq) is a powerful technique that enables the analysis of gene expression at the individual cell level. Bioinformatic tools for scRNA-seq data analysis have many different options throughout the typical scRNA-seq workflow (normalization, integration, annotation, clustering, and visualization), and the choice of method(s) and parameter(s) at each stage can impact results.

**Results:** Here, we introduce SWANS (v2.0), a configurable analysis pipeline that, in a single run, can employ multiple analysis methods, resolutions, and modifiable parameters. The resulting clustering arrangements, differential gene expression results, and other quantitative measurements can be dynamically visualized and compared in a Shiny interactive report to assist in choosing a single analysis schema for annotation and downstream analysis. Once a final approach is chosen, SWANS will perform differential gene expression (DGE) analysis based on experimental conditions and gene set enrichment analysis (GSEA) in addition to creating reports that display figures and interactive tables, quality control metrics, and benchmarking information. SWANS uses Snakemake as a workflow manager, Cell Ranger for alignment and gene expression quantification, Seurat for single cell data analysis, and additional single cell R packages for quality control and downstream single cell analysis.

**Conclusion:** SWANS is a tailorable pipeline that provides options for quality control, dimensionality reduction, clustering, differential gene expression analysis, gene set enrichment analysis, and trajectory analysis. Additionally, SWANS generates a series of reports that facilitate sharing large volumes of complex data in a clear and concise manner with other investigators.

## 1 Background

Advancements in single-cell and single-nucleus RNA-sequencing (scRNA-seq and snRNA-seq, respectively; scnRNA-seq used to refer to both methods collectively) have ushered forth an era concentrated on the transcriptional portraiture of individual cells. With scnRNA-seq analysis, researchers have gained insight into the heterogeneity in tissue composition, identified rare cell types, determined cell lineages, and advanced our understanding of biological processes [1–5]. Numerous tools and pipelines have been developed to aid researchers in managing and processing these data, and nearly all such tools have a variety of parameters and options [6–14]. The impact of selecting alternative parameters and/or approaches can alter cell cluster assignments, which further affects differential gene expression (DGE) analysis results, which in turn, influences the results of gene set enrichment analysis (GSEA) [15]. There is not a universal analysis approach that works for all scnRNA-seq data; tissue type, collection method, technique (single cell vs. single nucleus), the state of the tissue (*e*.*g*., formalin-fixed para”n-embedded, frozen, or fresh), sequencing depth, platform differences, and other factors can all impact the resulting sequencing data. This means that the optimal bioinformatic analysis approach may differ between datasets. However, it is not always clear which set of methods will be the most suitable for a particular dataset, and it can be valuable to compare approaches to determine which set of options produces results most reflective of the biological reality [16, 17].

Seurat (v5) [1, 3, 8, 9, 18] is a widely used tool for scRNA-seq analysis that offers multiple methods for normalization, scaling, and integration—steps that adjust for technical variation and aligns data across experimental conditions. These processed data are used to build a shared nearest-neighbor (SNN) graph, which represents relationships between cells based on their gene expression profiles, and is subsequently used to identify clusters of cells. Choices at each step (*e*.*g*., normalization, integration, the clustering granularity (resolution)) lead to different cluster outcomes, affecting downstream analysis and biological conclusions. Therefore, it is valuable to compare methods in Seurat to determine the most biologically realistic procedure, but performing these comparisons manually and compiling the results can be cumbersome.

There are numerous tools developed to work with Seurat that are not intrinsically part of its workflow. For example, Seurat takes in gene expression count data as input, which typically needs to be generated using a program like Cell Ranger [19] for read alignment and quantification of transcripts. Before analyzing Cell Ranger output in Seurat, investigators often preprocess the data using tools such as SoupX [20], which removes ambient RNA (free-floating transcripts that can be captured in droplets and confound downstream analysis), and DoubletFinder [21], which identifies and flags droplets containing multiple cells (‘doublets’) for exclusion [22]. Additionally, there are many tools for post-Seurat analysis such as fgsea, an R package for GSEA [23], Monocle3, an R package for trajectory analysis [24–27], and CellPhoneDB [28] and CellChat [29], R packages for intercellular communication analysis. Each tool comes with its own documentation and data formatting expectations, requiring users to appropriately structure Seurat-derived data to ensure compatibility, which can be a time-consuming process. Moreover, compiling and consolidating the resulting outputs into a coherent and interpretable form for effective communication also demands thoughtful visualization, and requires users to determine how best to organize, condense, and display numerous complex tables and figures in a way that is both informative and visually accessible without overwhelming the reader.

We introduce SWANS (Single-entity Workflow ANalysiS), a comprehensive framework for pre-processing and post-annotation analysis of sncRNA-seq data. SWANS leverages Seurat (v5.1.0) and Snakemake [16, 30] as a workflow manager and allows users the flexibility to control a wide range of analysis parameters through YAML configuration files [31]. Snakemake is a powerful framework for building scalable, reproducible, and e”cient data analysis pipelines. Its rule-based structure defines each step, its dependencies, and tools, ensuring clarity and reproducibility. By modifying YAML configuration files, users can easily adapt pipelines to new analyses without altering the underlying code. Snakemake also e”ciently resumes interrupted workflows, automatically handles parallelization, and tracks resource usage (*e*.*g*., memory, time), simplifying debugging and optimization making it a robust solution for managing both small and large-scale computational workflows. While existing scnRNA-seq pipelines typically emphasize raw data processing and gene count generation ([32, 33]), SWANS includes options for quality control, data normalization, dimensionality reduction, clustering, DGE analysis, GSEA, trajectory inference, and a wide range of visualizations and reports.

SWANS was developed with three key design features: 1. dynamic comparison of the many different combinations of methods and analysis operations available in Seurat in an interactive report; 2. seamless workflow between Cell Ranger, Seurat, and R packages used for pre-processing, quality control, and post-Seurat analysis of scRNA-seq data; and 3. presentation of data in a clear, concise, and simplified manner, enabling investigators to connect results to biological insights. To evaluate our workflow, we downloaded eight publicly available datasets with varied sample sizes, tissue states, and experimental conditions, then processed them through SWANS with various customizations. We selected one dataset to showcase all reports and interactive visualizations while also leveraging Snakemake’s benchmarking capabilities to evaluate each step of the analysis pipeline and demonstrate the time and memory resources used for each dataset.

## 2 Implementation

### 2.1 Pipeline overview

SWANS is split into two phases: the preliminary analysis and post-annotation analysis (Figure 1). The purpose of the preliminary analysis is to compare different clustering arrangements to find the optimal configuration and determine the cell types based on differentially expressed genes (DEGs) between clusters and other quantitative metrics for the selected configuration. The preliminary results can be visualized collectively in a Shiny app to allow the user to compare the various combinations of results [34]. Once an optimal configuration is chosen by the user, the post-annotation analysis can be executed and includes DGE analysis and GSEA/pathway analysis to compare experimental conditions within each cluster, and optionally pursue trajectory analysis.

**Fig. 1.**
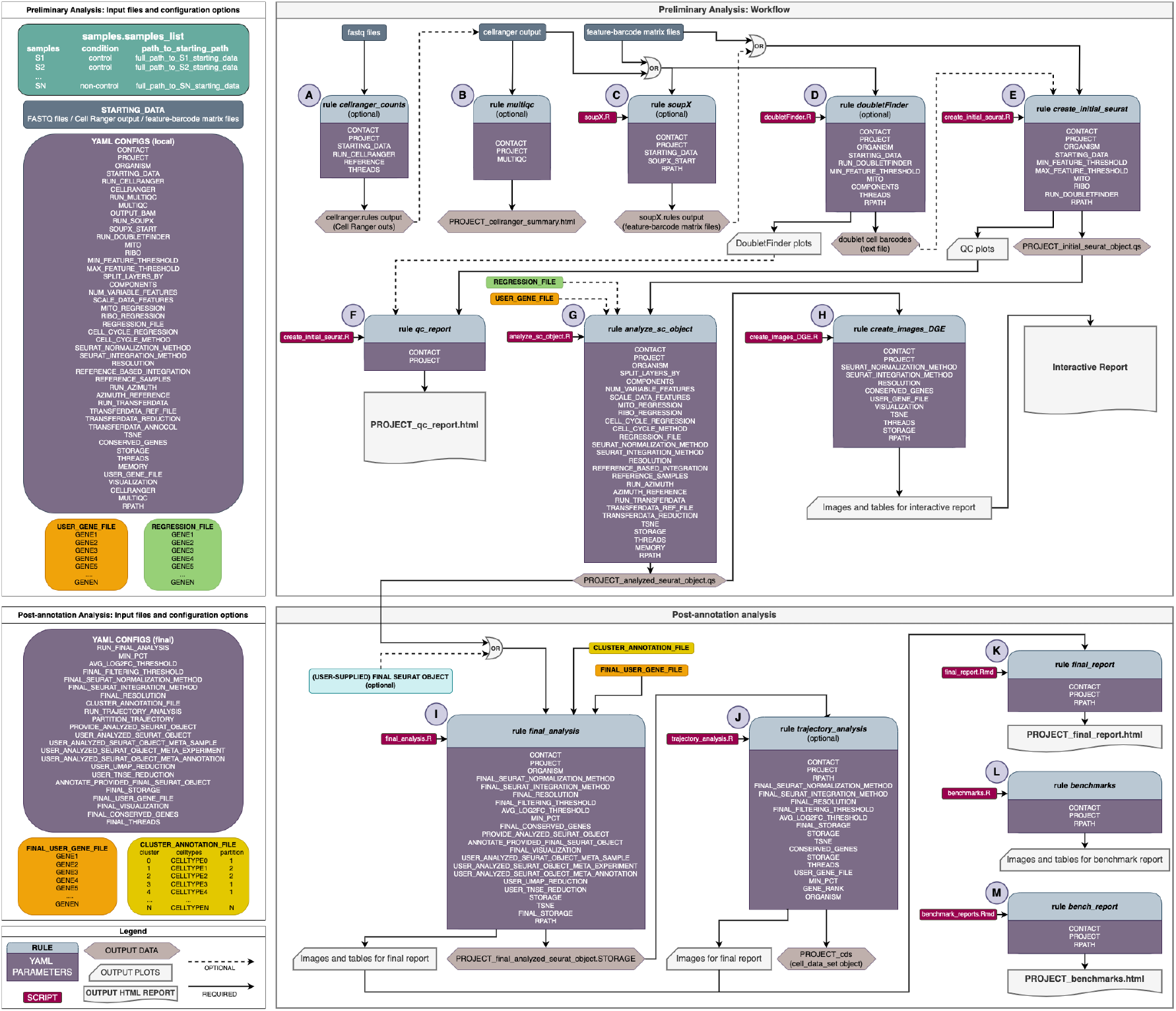
Schematic of SWANS Workflow. Preliminary analysis consists of steps A-H (configuration variables on upper left), the final analysis is included in I-M (configuration variables on lower left).

SWANS utilizes Snakemake as its workflow manager, enabling reproducibility, parallel computing, scalability, and e”cient management of computational resources. To run SWANS, the user must provide a file (`samples.sample_list`) containing the sample names, sample condition, and a path to the starting data. Snakemake supports the use of YAML configuration files, which define parameters that can be passed to specific Snakemake rules [31]. For each phase of SWANS (preliminary analysis and post-annotation analysis), there is a corresponding YAML configuration file that is used to specify parameters. The SWANS workflow consists of 14 Snakemake rules: nine for the preliminary analysis (Figure 1.A-H) and five for the post-annotation analysis (Figure 1.I-M).

#### 2.1.1 Rules: Preliminary Analysis

- *cellranger_counts* (optional): Starting with FASTQ files, the rule will align reads that are filtered using Cell Ranger’s `count` pipeline. Cell Ranger outputs filtered and raw feature-barcode matrices, secondary analysis results (e.g., clustering, PCA, t-distributed Stochastic Neighbor Embedding (t-SNE) [35] plots), quality control metrics, and an interactive HTML report summarizing the data.
- *multiqc_cellranger_report* (optional): This rule will generate a comprehensive report using MultiQC [36] which summarizes quality control metrics from multiple Cell Ranger outputs, including sequencing quality, read distribution, and gene expression metrics across samples.
- *soupX* (optional): This rule runs the tool SoupX, which corrects for ambient RNA contamination in single-cell RNA sequencing data to improve the accuracy of gene expression measurements. Outputs are corrected single-cell RNA sequencing count matrices with reduced ambient RNA contamination, along with diagnostic plots to assess the quality of the correction.
- *doubletFinder* (optional): Using DoubletFinder, this rule identifies singlets and doublets from single-cell RNA sequencing data and outputs a list of cells classified as doublets.
- *create_initial_seurat* : This rule filters cells from each sample based on user specifications (including doublets, minimum/maximum number of features per cell, %mito, %ribo), and merges all samples into a Seurat object.
- *make_qc_report* : This rule will create a report that provides an overview of employed QC tools and cell filtering parameters, predicts number of doublets/sample, metrics (number of cells, features, RNAfragments, mitochondria percentage, and ribosomal percentage) pre- and post-filtering of cells.
- *analyze_sc_object* : This rule performs optional gene regression, data normalization, cell cycle scoring, and PCA (via `RunPCA`). It also integrates the data, optionally annotates clusters using RunAzimuth and TransferData, calculates significant PCA components (with 90% variance), runs t-SNE (optional), identifies clusters with `FindClusters`, and generates a clustree plot.
- *memory* : This rule calculates memory requirements based on a cumulative number of cells and genes in each sample.
- *cluster_plots_DGE* : This rule creates Uniform Manifold Approximation and Projection (UMAP) [37, 38] plots and optionally t-SNE plots, and for each cluster, finds DEGs, conserved genes (optional), and z-score transformations. User-supplied genes are also visualized according to user specifications (optional).

#### 2.1.2 Rules: Post-Annotation Analysis

- *final_analysis*: This rule annotates clusters, and within each cluster, performs DGE analysis, finds conserved genes (optional), and calculates the z-score transformation based on experimental conditions. Several plots are also created (cell proportions, UMAP, t-SNE (optional) for individual clusters, cluster-experimental condition, and cluster-sample. Within cluster visualization of user-supplied genes across experimental conditions are also created.
- *trajectory_analysis*: (optional) This rule performs trajectory analysis using Monocle3 and outputs figures of the trajectory graph overlaying the annotated UMAP.
- *final_report* : This rule creates a summary report that contains images and tables from final analysis rule.
- *benchmarks*: This rule creates images and a table of time and memory resources used for all analysis steps.
- *bench_report* : This rule summarizes all output from the benchmark rule in an HTML report.

### 2.2 YAML parameters and configurations in SWANS

To customize SWANS, two YAML configuration files are provided: local configs.yaml for the preliminary analysis and final configs.yaml for the post-annotation analysis, each corresponding to the specific rules applied at their respective stages. All SWANS options are defined in the YAML files.

#### 2.2.1 Phase I: Preliminary analysis

The following items are the parameters available for customization in the preliminary analysis configuration file (local configs.yaml):

- *contact* : email will be sent when jobs complete
- *PROJECT* : project name (name of output directory)
- *ORGANISM* : organism (e.g., mouse, human)
- *STARTING_DATA*: starting point data (e.g., fastq, cellranger, matrix)
- *RUN_CELLRANGER*: run Cell Ranger count pipeline (y/n)
- *RUN_MULTIQC* : run multiQC on Cell Ranger output (y/n)
- *OUTPUT_BAM* : create bam file when running cellranger (y/n)
- *RUN_SOUPX* : run SoupX (y/n)
- *SOUPX_START* : starting files for SoupX: (outs, no clusters, h5)
- *RUN_DOUBLETFINDER*: run DoubletFinder (y/n)
- *MITO* : mito cutoff threshold
- *RIBO* : ribo cutoff threshold
- *MIN_FEATURE_THRESHOLD* : minimum feature threshold
- *MAX_FEATURE_THRESHOLD* : maximum feature threshold
- *SPLIT_LAYERS_BY* : metadata by which to split the object into layers (Experiment, Sample)
- *COMPONENTS* : number of Principal Components to use in Seurat
- *NUM_VARIABLE_FEATURES* : number of variable features
- *SCALE_DATA_FEATURES* : which features to use for data scaling (all or variable)
- *MITO_REGRESSION* : should mito percent be regressed? (y/n)
- *RIBO_REGRESSION* : should ribo percent be regressed? (y/n)
- *REGRESSION_FILE* : file of additional genes to be regressed
- *CELL_CYCLE_REGRESSION* : should cell cycle be regressed? (y/n)
- *CELL_CYCLE_METHOD* : which cell cycling method (standard, alternative)
- *SEURAT_NORMALIZATION_METHOD* : normalization method for Seurat (sct, standard)
- *SEURAT_INTEGRATION_METHOD* : integration method for Seurat (cca, harmony, rpca)

- *REFERENCE BASED INTEGRATION* : run integration in reference-based fashion? (y/n)
- *REFERENCE SAMPLES* : which sample(s) to use as references for ref-based integration?
- *RUN_AZIMUTH* : run Azimuth annotation in Seurat? (y/n)
- *AZIMUTH REFERENCE* : human ref options: adiposeref, bonemarrowref, fetusref, heartref, humancortexref, kidneyref, lungref, pancreasref, pbmcref, tonsilref; mouse ref options: mousecortexref
- *RUN_TRANSFERDATA*: run TransferData in Seurat to annotate with a provided reference Seurat object? (y/n)
- *TRANSFERDATA_REF_FILE* : path to reference Seurat object for TransferData? (path/to/seuratfile.storage)
- *TRANSFERDATA_REDUCTION* : which dimensional reduction in the TRANS-FERDATA REF FILE to use for TransferData?
- *TRANSFERDATA_ANNOCOL*: which Seurat meta.data column to use for TransferData annotation?
- *RESOLUTION* : resolution value(s) examples: single value: 0.5; multiple values: 0.1,0.3,0.5
- *TSNE* : make t-SNE plots (y/n)
- *CONSERVED_GENES* : run conserved genes (y/n)
- *STORAGE* : additional storage format (rds,scEO)
- *THREADS* : number of threads
- *MEMORY* : max memory
- *USER_GENE_FILE* : full path with filename to file containing genes of interest (optional)
- *VISUALIZATION* : which visualization for USER GENE FILE (feature,violin,ridge,dot) – choose 1 or more

There are three different options for the format of input data in SWANS: FASTQ files, Cell Ranger ‘outs’ files, or files in the sparse matrix Market Exchange (MEX) format (feature-barcode matrix files). There is also the option to create BAM files while running Cell Ranger, and for running a MultiQC report on the generated or supplied Cell Ranger files in the ‘outs’ directory.

The preliminary analysis includes optional quality control tools that can be run on each sample to address potential artifacts. Two artifacts of droplet-based scnRNA-seq experiments that contribute to batch effects are the introduction of non-biological ambient mRNA, and multiple cells captured by a single bead but interpreted as a single cell (called “doublets” or “multiplets”). The user can optionally choose to run SoupX to remove background-contaminating ambient RNA, and DoubletFinder to identify and remove multi-cellular droplets presenting as a single cell.

Thresholds are applied to filter out cells with an excessive or insu”cient number of features, or those with disproportionately high expression of mitochondrial or ribosomal genes. Individual Seurat objects are created for each sample and filtered according to these user-defined thresholds. All Seurat objects are then merged into one object for normalization. Users can opt to regress out mitochondrial, ribosomal, and cell cycle genes, or provide a file with specific genes to be scored for regression. After normalization, the pipeline performs dimensionality reduction, integration, DGE analysis for each cluster, and visualization.

The user can choose how many initial principal components to explore, set the number of variable genes, and control which genes are used to scale the analysis (variable or all).

The user can define which normalization method(s) and integration method(s) should be performed. Normalization options include the Seurat “standard approach” for normalization (`NormalizeData`, `FindVariableFeatures`, and `ScaleData` work-flow) and the “sctransform” method (`SCTransform` function). An overview of the key differences between the Standard Approach and SCTransform, particularly in terms of normalization methods, feature selection, and output is provided in Table 1. Principal component analysis (PCA, implemented via `RunPCA` in Seurat) will be calculated for whichever of the normalization methods are run. Integration is performed using the `IntegrateLayers` function in Seurat and will be run for each normalization method and PCA reduction. Any or all of the following integration methods can be included in the analysis: `CCAIntegration` (canonical correlation analysis (CCA) [9], `HarmonyIntegration` (Harmony method [39], and `RPCAIntegration` (reciprocal PCA (RPCA) [40]). Table 2 provides a comparison of the three integration methods—CCA, Harmony, and RPCA — highlighting key aspects such as method type, batch effect handling, and output characteristics.

**Table 1.**
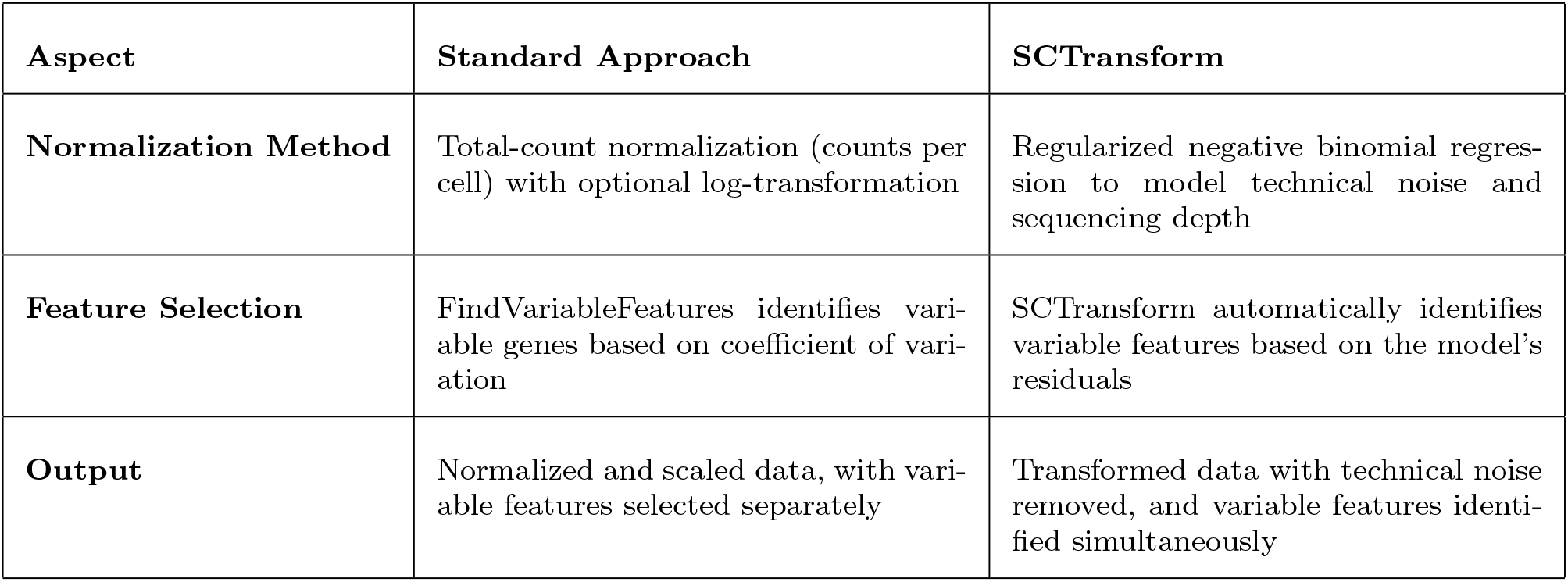
Normalization Comparison of Standard Approach and SCTransform

| Aspect | Standard Approach | SCTransform |
| --- | --- | --- |
| Normalization Method | Total-count normalization (counts per cell) with optional log-transformation | Regularized negative binomial regression to model technical noise and sequencing depth |
| Feature Selection | FindVariableFeatures identifies variable genes based on coefficient of variation | SCTransform automatically identifies variable features based on the model's residuals |
| Output | Normalized and scaled data, with variable features selected separately | Transformed data with technical noise removed, and variable features identified simultaneously |

**Table 2.**
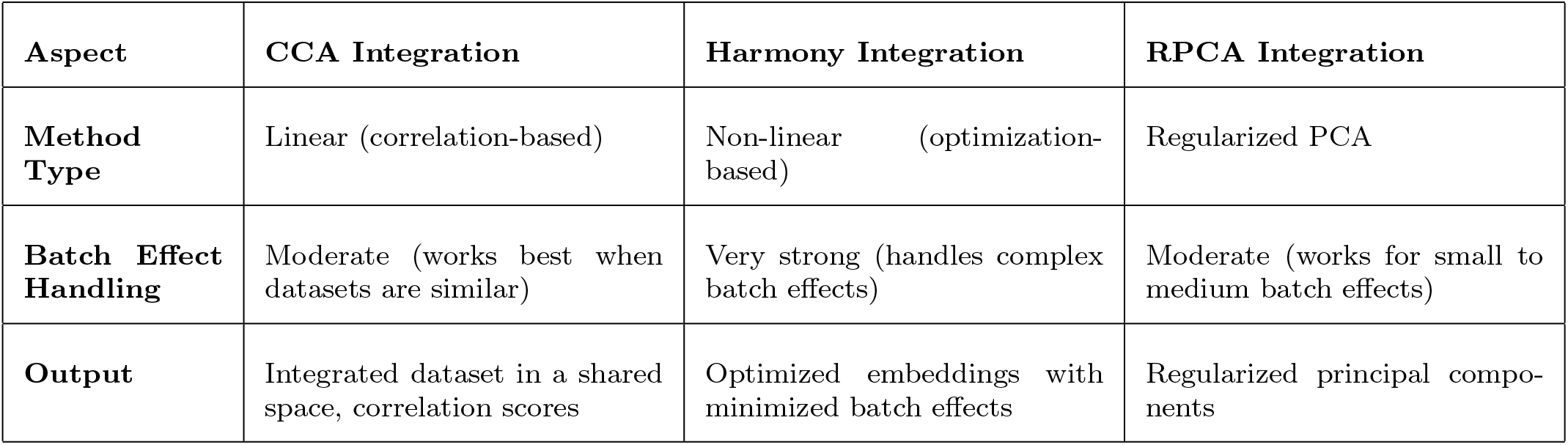
Comparison of Integration Methods: CCA, Harmony, and RPCA

| Aspect | CCA Integration | Harmony Integration | RPCA Integration |
| --- | --- | --- | --- |
| Method Type | Linear (correlation-based) | Non-linear (optimization-based) | Regularized PCA |
| Batch Effect Handling | Moderate (works best when datasets are similar) | Very strong (handles complex batch effects) | Moderate (works for small to medium batch effects) |
| Output | Integrated dataset in a shared space, correlation scores | Optimized embeddings with minimized batch effects | Regularized principal components |

The user can select any combination of approaches which will be uniquely stored within the Seurat object. For example, if the user chooses to run both normalization methods (by specifying in the SWANS configuration file `standard` for the Seurat standard approach and `sct` for Seurat’s “sctransform” method), and all three integration approaches (`cca` for `CCAIntegration`, `harmony` for `HarmonyIntegration`, and `rpca` for `RPCAIntegration`), at resolutions 0.1, 0.2, and 0.3, the resulting Seurat object will have clustering results in the object metadata for all 18 possible combinations of what we call ‘analysis schema’. By calculating all of the possibilities in a single run of SWANS, the user can view all of the normalization, integration, and clustering combinations (schemas) in the Shiny interactive report and select the most suitable schema for their data. Users can also choose to run reference-based integration and select which samples will be used as the reference for integration, which is helpful when integrating large datasets.

Conserved genes exhibit consistent expression patterns across different conditions in clusters, indicating their potential importance in biological processes. The user can opt to run Seurat’s `FindConservedMarkers` to find conserved genes and they may also choose to include t-SNE figures in addition to the UMAP figures that are produced by default. Users also have the option of supplying a text file of markers genes they would like to have visualized on feature, dot, ridge, and/or violin plots. If there are multiple resolutions specified in the configuration file, the R package clustree [41] will be run to produce a clustering tree image that can be used to compare cell assignment (to clusters) across resolutions.

For annotation, Azimuth (via `RunAzimuth` from the Azimuth R package [42]) can be used to identify cell types based on the Azimuth references that are included in the SeuratData package [43]), or they can use Seurat’s `TransferData` function to transfer the annotations from a provided Seurat object to the current working dataset.

This workflow also takes advantage of the future framework R package [44] to speed up Seurat commands and if the user is working on a system with multiple cores, they can set the number of threads (number of central processing unit (CPU) cores) that can be used for parallel processing.

Once the preliminary analysis is complete, a QC report and an interactive report (Shiny application) are generated. The QC report provides a comprehensive overview of the data quality at both the initial stage and after cell filtering. It helps users assess the integrity and reliability of the raw data, as well as the impact of cell filtering on the dataset, ensuring that only high-quality data is retained for downstream analysis. The Shiny app displays the interactive report and can be used to dynamically view the results of the SWANS preliminary analysis phase and allows the user to easily compare multiple analysis schemas side-by-side. The user can dynamically inspect and compare multiple clustering resolutions visualized on UMAPs, alongside the related DEGs, z-scores, and cell counts/proportions for each cluster therein. During the preliminary analysis, Clustree will also be run, which allows the user to see how cells move as a function of resolution. All of this information should be used to determine the optimal combination of options for downstream analysis. In addition to the interactive report, a PDF will be created for each of the analysis schemas. Each PDF includes four UMAP images (colored by clusters, split by experimental conditions, split by samples, colored by cell cycling phase), bar plots of cell proportions (by sample), and visualization plots for any user-supplied genes. Each PDF is paired with 3 tables (cell counts per cluster (by sample), DEG results (by cluster), top 100 DEGs (by cluster) and an optional table of conserved markers (by cluster) if requested by user.

#### 2.2.2 Phase II: Post-Annotation analysis

After the user has reviewed the results of the preliminary analysis, they can choose their preferred analysis schema and indicate how they would like to annotate the cell types in their dataset for the post-annotation analysis. The post-annotation analysis will use the Seurat file output from the preliminary analysis. The following parameters are available for customization in the post-annotation analysis configuration file (final configs.yaml):

- *RUN_FINAL_ANALYSIS* : run post-annotation analysis (y/n)
- *MIN_PCT* : minimum % of cells expressing gene in either compared groups for DGE
- *AVG_LOG2FC_THRESHOLD* : filtering DEGs (avg log2FC) – single value
- *FINAL_FILTERING_THRESHOLD* : filtering threshold/value of adj.p.value for pathway/GSEA results – single value
- *FINAL_SEURAT_NORMALIZATION_METHOD* : final normalization method for Seurat (standard,sct)
- *FINAL_SEURAT_INTEGRATION_METHOD* : final integration method for Seurat (cca,rpca,harmony)
- *FINAL_RESOLUTION* : final resolution value – single value
- *CLUSTER_ANNOTATION_FILE* : file with names of new clusters
- *RUN_TRAJECTORY_ANALYSIS* : run Monocle3 trajectory analysis? (y/n) – optional
- *PARTITION_TRAJECTORY* : partition clusters (y/n)
- *PROVIDE_ANALYZED_SEURAT_OBJECT* : supply final Seurat object (y/n)
- *USER_ANALYZED_SEURAT_OBJECT* : path to Seurat object – cannot be blank if PROVIDE ANALYZED SEURAT OBJECT = ‘y’
- *USER_ANALYZED_SEURAT_OBJECT_META_SAMPLE* : meta data character string to access ‘Sample’ in Seurat object (e.g., Samples, sample name’)

- *USER_ANALYZED_SEURAT_OBJECT_META_EXPERIMENT* : meta data character string to access ‘Experiment’ in Seurat object (e.g., Experiment, Conditions)
- *USER_ANALYZED_SEURAT_OBJECT_META_ANNOTATION* : meta data character string that holds annotation information in Seurat object (e.g., celltypes, annotation layer)
- *USER_UMAP_REDUCTION* : UMAP reduction in supplied Seurat object (e.g., standard.cca.umap)
- *USER_TNSE_REDUCTION* : t-SNE reduction in supplied Seurat object (e.g., standard.rpca.tsne)
- *ANNOTATE_PROVIDED_FINAL_SEURAT_OBJECT* : annotate Seurat object? (y/n) (only if providing Seurat object)
- *FINAL_STORAGE* : additional final storage format (qs,rds,sceo,cloupe,cellchat,cellphone)
- *FINAL_USER_GENE_FILE* : full path with filename to file containing genes of interest
- *FINAL_VISUALIZATION* : which visualization(s) for FINAL USER GENE FILE (feature,violin,ridge,dot)
- *FINAL_CONSERVED_GENES* : run conserved genes (y/n)
- *FINAL_THREADS* : number of threads

The final resolution and methods for normalization and integration can be specified in the post-annotation configuration YAML file. The user must also provide a tab-delimited text file that indicates how the clusters should be annotated. For each cluster, DEGs are found by comparing transcriptional differences between experimental conditions. Using the Molecular Signatures Database (MSigDB), the fgsea R package [23, 45], and the differentially expressed genes, gene set enrichment analysis is performed on for each cluster. In SWANS, GSEA is performed for two different ranking metrics (average log2 fold change and adjusted p-value) and for three of the gene set collections available from MSigDB (hallmark, canonical pathways, and gene ontology).

Additionally, z-score transformation of transcription gene counts and cell counts/proportions are calculated to characterize each cluster, each experimental condition within each cluster, and each sample within each cluster. The z-score transformation standardizes gene expression data by converting them to a common scale. This removes biases due to differences in scale making it easier to compare transcript profiles between experimental conditions and samples within a cluster. The user can specify whether to find conserved genes for each cluster, and the user can also provide a list of genes for visualization, and define how they would like those genes visualized (feature, dot, ridge, and/or violin plots). As with the local configs.yaml, the user can specify the number of threads for parallel processing, and the final format of their data. For post-annotation analysis, there are additional options to output a `SingleCellExperiment` object (via the SingleCellExperiment package [46]), `.cloupe` files for viewing in the 10X Genomics Loupe Browser (v8.1.0) [47], and/or a `CellChat` object from the R package CellChat to infer intercellular communication. Although time consuming, the user can also write the normalized gene expression data as a matrix file to infer intercellular communication with `cellphoneDB`. Finally, there is an option to perform trajectory analysis using the Monocle 3 R package. The results from the post-annotation analysis will be compiled into an HTML report so that the user can easily view and share the document with collaborators. At the conclusion of the post-annotation analysis, the benchmarking information output by Snakemake is compiled into a comprehensive HTML report to provide users with insights into the time and resources allocated to each step (rule) of the pipeline.

## 5 Results

### 5.1 Preliminary Analysis Output

To demonstrate the options available in SWANS and its overall usage, we downloaded eight datasets, one from the ArrayExpress database (E-MTAB-13067) [48], two from 10X genomics (human frozen bone marrow [49–53], brain cells from an E18 Mouse [54–59]), and five from NCBI (PRJNA1010957 (GEO: GSE242001), PRJNA1095970 (GEO:GSE263228), PRJNA1006693 (GEO: GSE241184), PRJNA1185392 (GEO: GSE281736), PRJNA790856 (GEO: GSE191288)) [60–64]), and processed them with SWANS for benchmarking purposes.

To demonstrate specific YAML configurations and to provide examples of all reports generated by SWANS, we used a dataset of seven thyroid samples from PRJNA790856 (GEO: GSE191288). The GSE191288 dataset is related to research that investigated the tumor microenvironment (TME) in bilateral papillary thyroid carcinoma (PTC) through analysis of single-cell RNA sequencing of cells from three bilateral PTC pairs and one non-tumor thyroid sample [65]. The GEO data was downloaded in H5 format and was converted to MEX formatted files using the Bioconductor [66] package DropletUtils [67, 68]. The `samples.sample_list` file (Table 3) includes the sample names, conditions, and paths to the folders containing the MEX files.

**Table 3.**
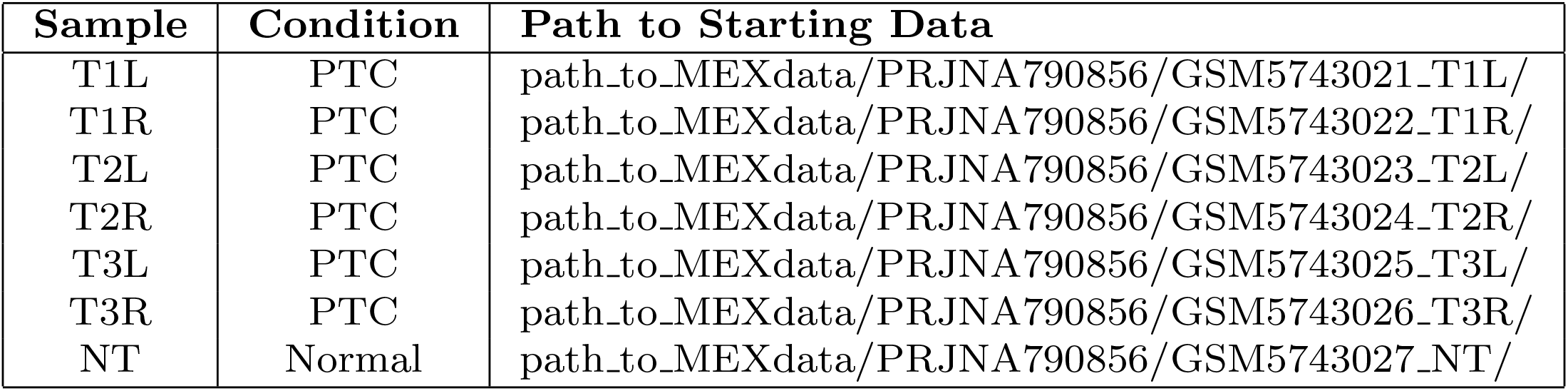
Example of the `samples.sample_list` file, which includes sample information — sample name, condition, and the corresponding paths to directories containing MEX files.

| Sample | Condition | Path to Starting Data |
| --- | --- | --- |
| T1L | PTC | path_to_MEXdata/PRJNA790856/GSM5743021_T1L/ |
| T1R | PTC | path_to_MEXdata/PRJNA790856/GSM5743022_T1R/ |
| T2L | PTC | path_to_MEXdata/PRJNA790856/GSM5743023_T2L/ |
| T2R | PTC | path_to_MEXdata/PRJNA790856/GSM5743024_T2R/ |
| T3L | PTC | path_to_MEXdata/PRJNA790856/GSM5743025_T3L/ |
| T3R | PTC | path_to_MEXdata/PRJNA790856/GSM5743026_T3R/ |
| NT | Normal | path_to_MEXdata/PRJNA790856/GSM5743027_NT/ |

The YAML parameters for the preliminary analysis of GSE191288 are listed in Table 4. The GSE191288 dataset was converted to feature-barcode matrix files which contain the gene counts, so Cell Ranger was not needed for alignment and quantification. In addition, MultiQC and SoupX were not used in this case because those tools require data from Cell Ranger outs files, which were not available for this dataset. However, DoubletFinder was employed to identify and remove cells identified as doublets. SWANS filters each created Seurat object according to the thresholds set in the configuration file; in this case, SWANS filtered the data to retain cells with more than 200 features (genes), less than 4000 features, and with no more than 20% mitochondrial gene expression.

**Table 4.** YAML Parameters used in the preliminary analysis of PRJNA790856 (GEO: GSE191288).

|  |  |
| --- | --- |
| **contact** |  |
| **PROJECT** | prjna790856 |
| **ORGANISM** | human |
| **STARTING_DATA** | matrix |
| **RUN_CELLRANGER** | No |
| **RUN_MULTIQC** | No |
| **OUTPUT_BAM** | No |
| **RUN_SOUPX** | No |
| **SOUPX_START** | no_clusters |
| **RUN_DOUBLETFINDER** | Yes |
| **MITO** | 20 |
| **RIBO** | 100 |
| **MIN_FEATURE_THRESHOLD** | 200 |
| **MAX_FEATURE_THRESHOLD** | 4000 |
| **SPLIT_LAYERS_BY** | Sample |
| **COMPONENTS** | 30 |
| **NUM_VARIABLE_FEATURES** | 3000 |
| **SCALE_DATA_FEATURES** | variable |
| **MITO_REGRESSION** | No |
| **RIBO_REGRESSION** | No |
| **REGRESSION_FILE** | NA |
| **CELL_CYCLE_REGRESSION** | No |
| **CELL_CYCLE_METHOD** | standard |
| **SEURAT_NORMALIZATION_METHOD** | sct,standard |
| **SEURAT_INTEGRATION_METHOD** | cca,harmony,rpca |
| **REFERENCE_BASED_INTEGRATION** | No |
| **REFERENCE_SAMPLES** | NA |
| **RUN_AZIMUTH** | No |
| **AZIMUTH_REFERENCE** | NA |
| **RUN_TRANSFERDATA** | No |
| **TRANSFERDATA_REF_FILE** | NA |
| **TRANSFERDATA_REDUCTION** | NA |
| **TRANSFERDATA_ANNOCOL** | NA |
| **RESOLUTION** | 0.1,0.2,0.3 |
| **TSNE** | No |
| **CONSERVED_GENES** | No |
| **STORAGE** | NA |
| **THREADS** | 30 |
| **MEMORY** | 1.38906E+12 |
| **USER_GENE_FILE** | prjna1185392.txt |
| **VISUALIZATION** | dot |

#### 5.1.1 QC Report

The first generated report provides a comprehensive review of all QC metrics, summarized in Figure 3. The QC report includes UMAP plots with cells color-coded according to their classification as either doublets or singlets (if DoubletFinder was run), as well as a violin plot illustrating the distribution of cell features for each classification, the total number of cells pre- and post-doublet removal, and the doublet rate (Figure 3.B). The QC report also displays violin plots with the number of features (nFeatures RNA), number of unique molecular identifiers (UMIs) (nCount RNA), mitochondrial percentage (percent.mito) and ribosomal percentage (percent.ribo) before and after filtering (Figure 3.C), and un-filtered and filtered Seurat FeatureScatter plots for combinations of “nCount_RNA”, “percent.mt”, “percent.ribo”. “nCount_RNA”, and “nFeature_RNA” (Figure 3.D-E). The full report includes YAML QC parameters, and DoubletFinder classification figures (*e*.*g*., Figure 3.B) for each sample in the dataset. A full version of the HTML report in PDF format is located in Supplementary materials (Supplementary 1).

#### 5.1.2 Interactive Report

To illustrate the utility of the interactive report, SWANS was run on the GSE191288 dataset using both normalization methods and all three integration approaches at three resolutions: 0.1, 0.2, and 0.3. This resulted in 18 unique analysis schemas (i.e., combination of normalization method, integration method, and resolution (Figure 2)), each with its own UMAP projection and a”liated metrics. The Shiny app has drop-down menus that can be used to select three different schemas for side-by-side comparison, and each schema’s UMAP visualization is dynamically displayed (Figure 4A.). Using the drop-down menus, the user can opt to show either the differentially expressed genes, z-scores of gene expression, or cluster cell counts and proportions for the displayed/selected schema above (Figure 4B.). Because there were multiple (*>*1) clustering resolutions requested in the configuration file in this example, clustree images were generated and reflect the cell assignment to clusters as a function of resolution (Figure 4C.). As a final note on z-score transformations, they are included in the interactive report to address specific cases where a single anatomical body contains multiple cell subtypes that are split into distinct clusters. In such cases, DGE analysis compares one subtype against others within the same body, which may not yield informative DEGs for annotation purposes.

**Fig. 2.**
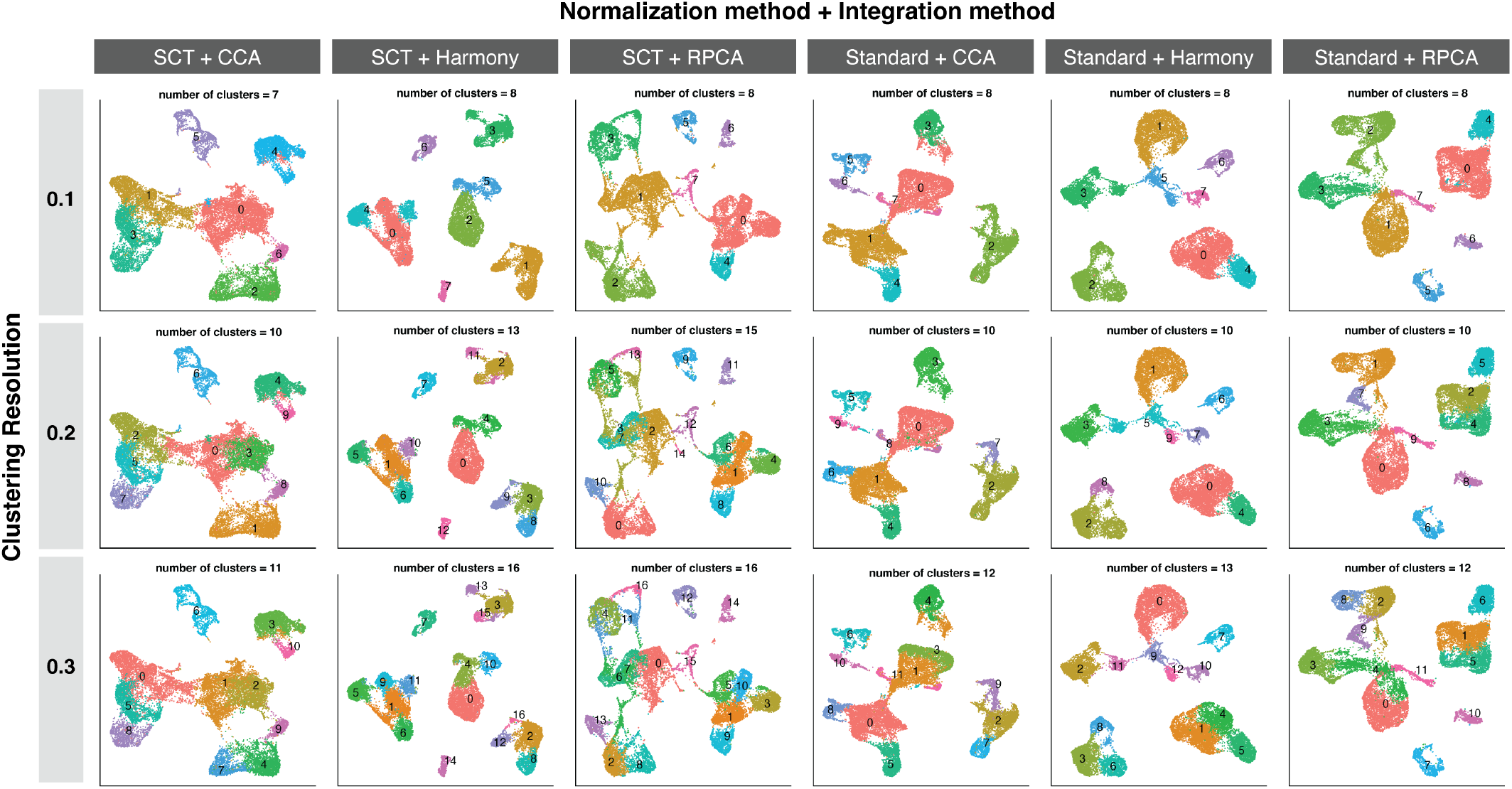
UMAP images displaying clusters for the 18 analysis schemas that result from combining each of Seurat’s two normalization methods (‘Standard Approach’ and SCTransform) with three of Seurat’s integration methods (CCA, Harmony, and RPCA) at three clustering resolutions (0.1, 0.2, and 0.3). Above each UMAP is the number of clusters for that combination of methods (x-axis: Normalization method + Integration method) at the specified clustering resolution (y-axis: Clustering Resolution).

**Fig. 3.**
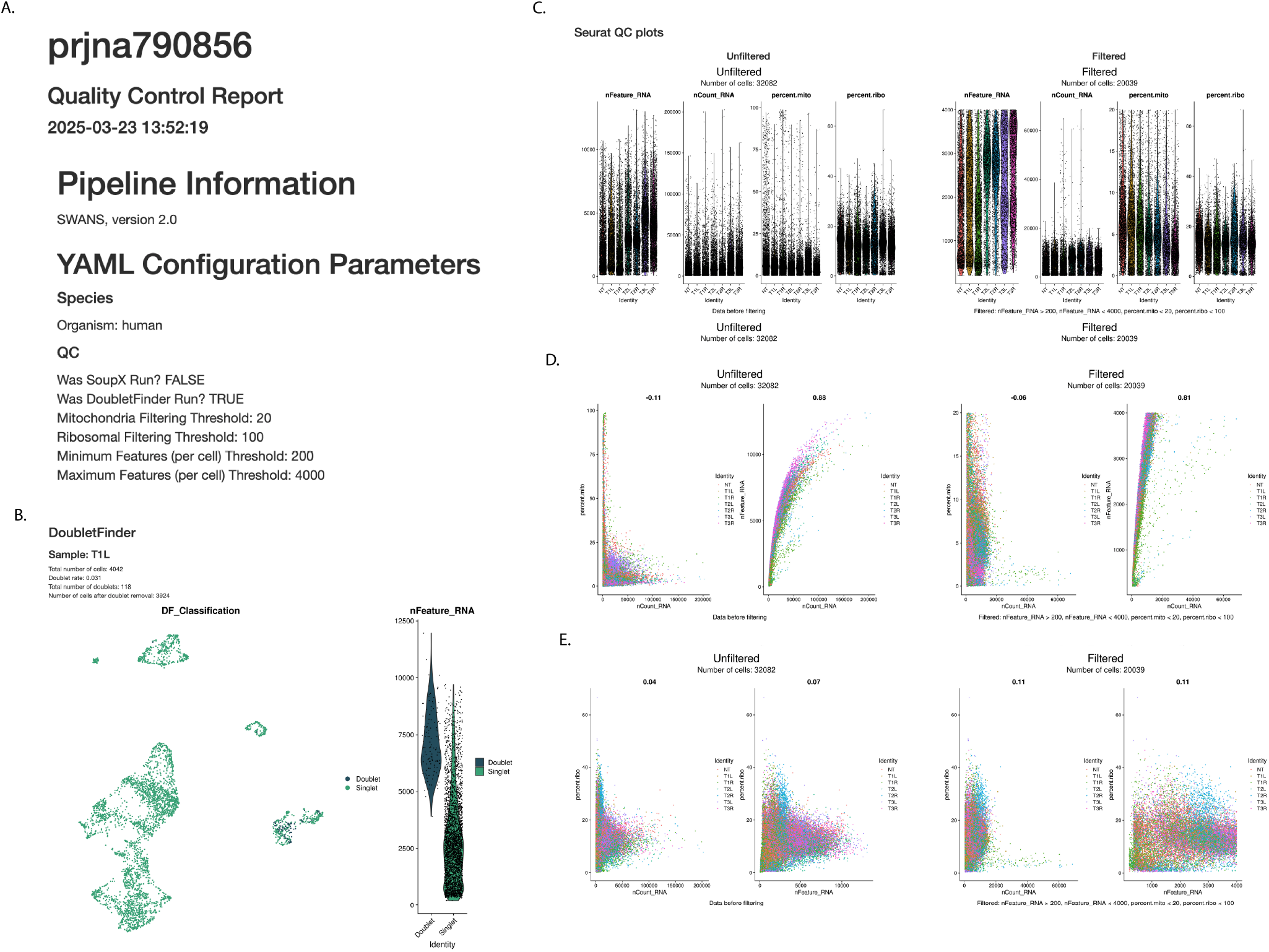
Excerpts from a QC report. A. Overview of employed QC tools and cell filtering parameters. B. Total number of cells, doublet rate, and number of doublets identified for a sample in the dataset (full report shows DoubletFinder results for each sample in dataset). C. Unfiltered (left) and filtered (right) number of features, RNA fragments, mitochondrial percentage, and ribosomal percentage in each cell across all samples in dataset. D. Per cell unfiltered (left) and filtered (right) feature scatter plots (number of UMIs vs. mitochondrial percentage and number of RNA fragments vs number of genes) colored by sample. E. Per cell unfiltered (left) and filtered (right) feature scatter plots (number of RNA fragments vs. ribosomal percentage and number of genes vs. ribosomal percentage), colored by sample. The full report is interactive and in HTML format; an example of this report in PDF format can be found in Supplementary 1.

**Fig. 4.**
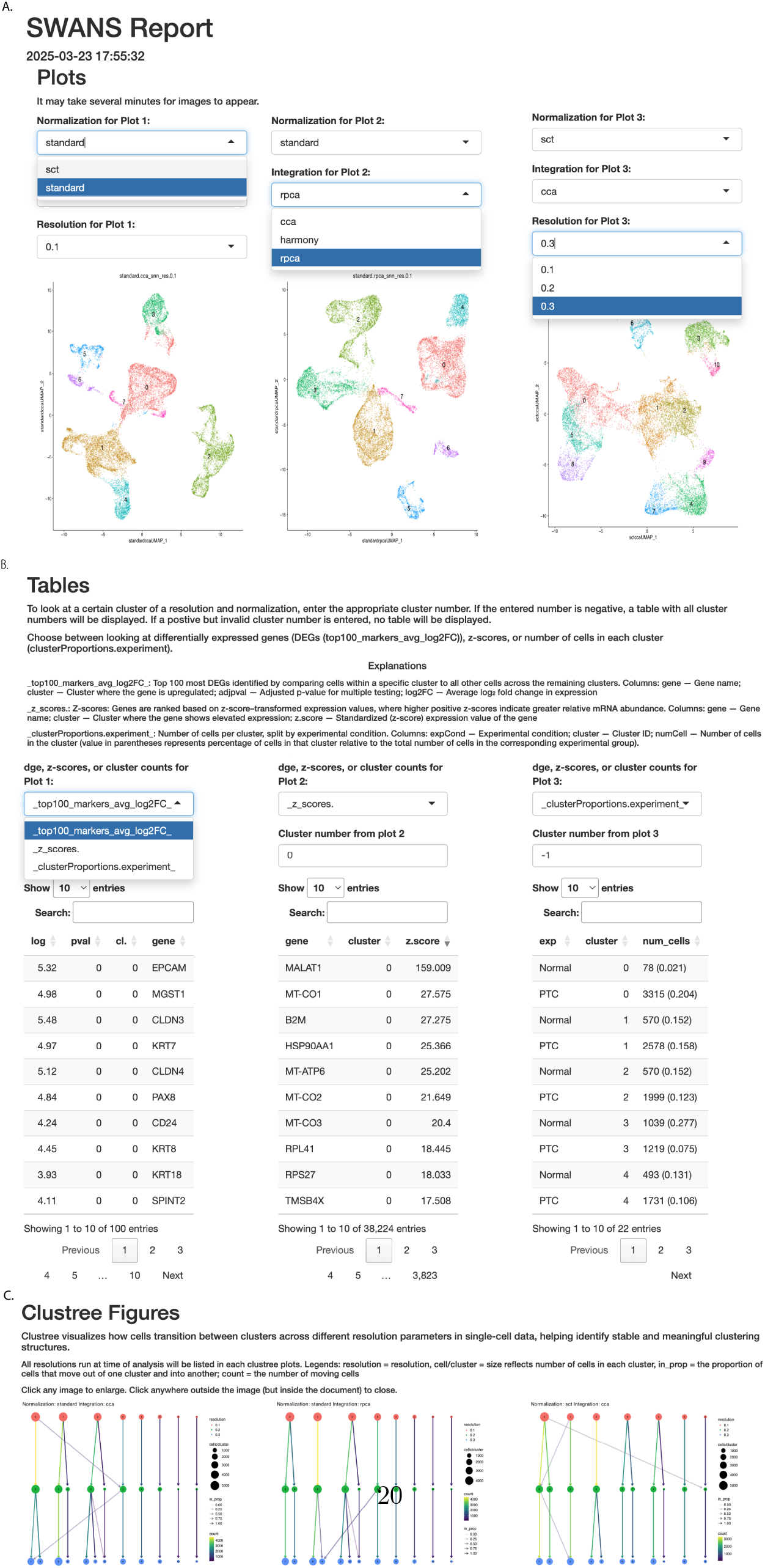
Shiny Interactive Report. A. Up to three (3) clustering schema can be dynamically selected from the menu for normalization, integration, and resolution. B. Table selection for top 100 DEGs, z-scores, and the proportion of cells, all by cluster. Tables are updated to reflect the clustering schema chosen (as shown in A). C. Clustree images that show how cells are assigned to clusters as a function of resolution. Images are dynamically linked to the options chosen in part A.

#### 5.1.3 Additional Output

To assist with annotation, the following annotation markers were listed in the `USER\_GENE\_FILE`: (Myeloid) *LYZ, S100A8, S100A9, CD14* ; (T and NK) *CD3D, CD3E, CD3G, CD247* ; (B cell) *CD79A, CD79B, IGHM, IGHD* ; (Thyrocytes) *TG, EPCAM, KRT18, KRT19* ; (Fibroblasts) *COL1A1, COL1A2, COL3A1, ACTA2* ; (Endothelial) *PECAM1, CD34, CDH5, VWF*, and were visualized with dot plots. These plots are included in a PDF that is created for every combination of normalization and integration methods at each resolution. An overview of the resulting preliminary analysis PDF can be seen for standard normalization with RPCA integration at resolution 0.1 in Figure 5. A full version of the report is located in Supplementary PDF materials 2.

**Fig. 5.**
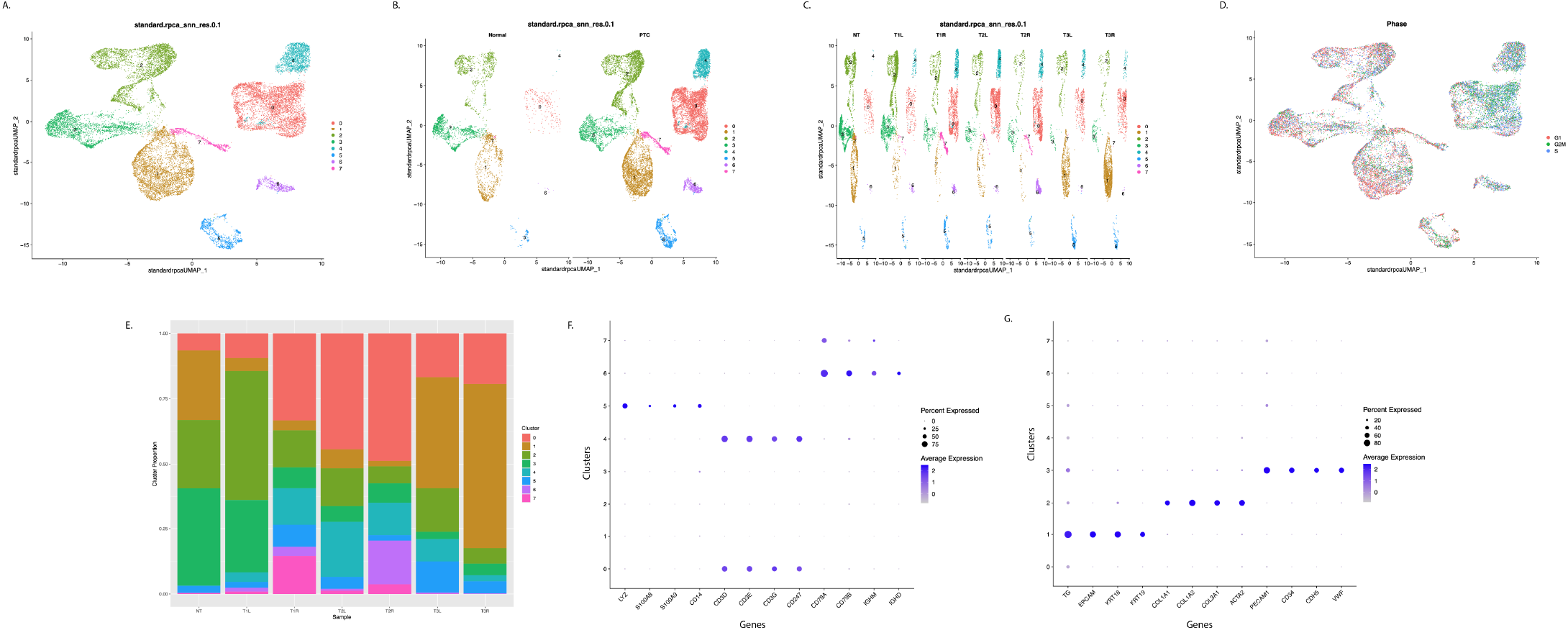
Example of additional images included in a PDF. Images include a composite UMAP colored by cluster (A), a UMAP split by experimental condition (B), a UMAP split by samples (C), and a UMAP colored by cell cycling phase (D). A barplot (E) is included to show the proportion of cells in each cluster for each sample, and images visualizing user-supplied genes in the format (dot, ridge, violin, and or feature plots) dictated in local configs.yaml file (F-G) (a maximum of 12 genes per image will be created). Images reflect standard normalization, rpca integration, at a resolution of 0.1. A PDF is created for all possible normalization, integration, and resolution combinations. A PDF rendering can be found in Supplementary 2.

### 5.2 Post-Annotation Analysis

The parameters for post-annotation analysis were defined for this dataset (Table 5). Using the aforementioned annotation markers, the most appropriate analysis schema was selected: standard normalization, RPCA integration, and a clustering resolution of 0.1 (`standard.rpca_snn_res.0.1`). Cluster identities were determined by manually examining the markers genes in the preliminary analysis, and the annotations for each cluster were defined in the `CLUSTER_ANNOTATION_FILE` (Table 6).

**Table 5.**
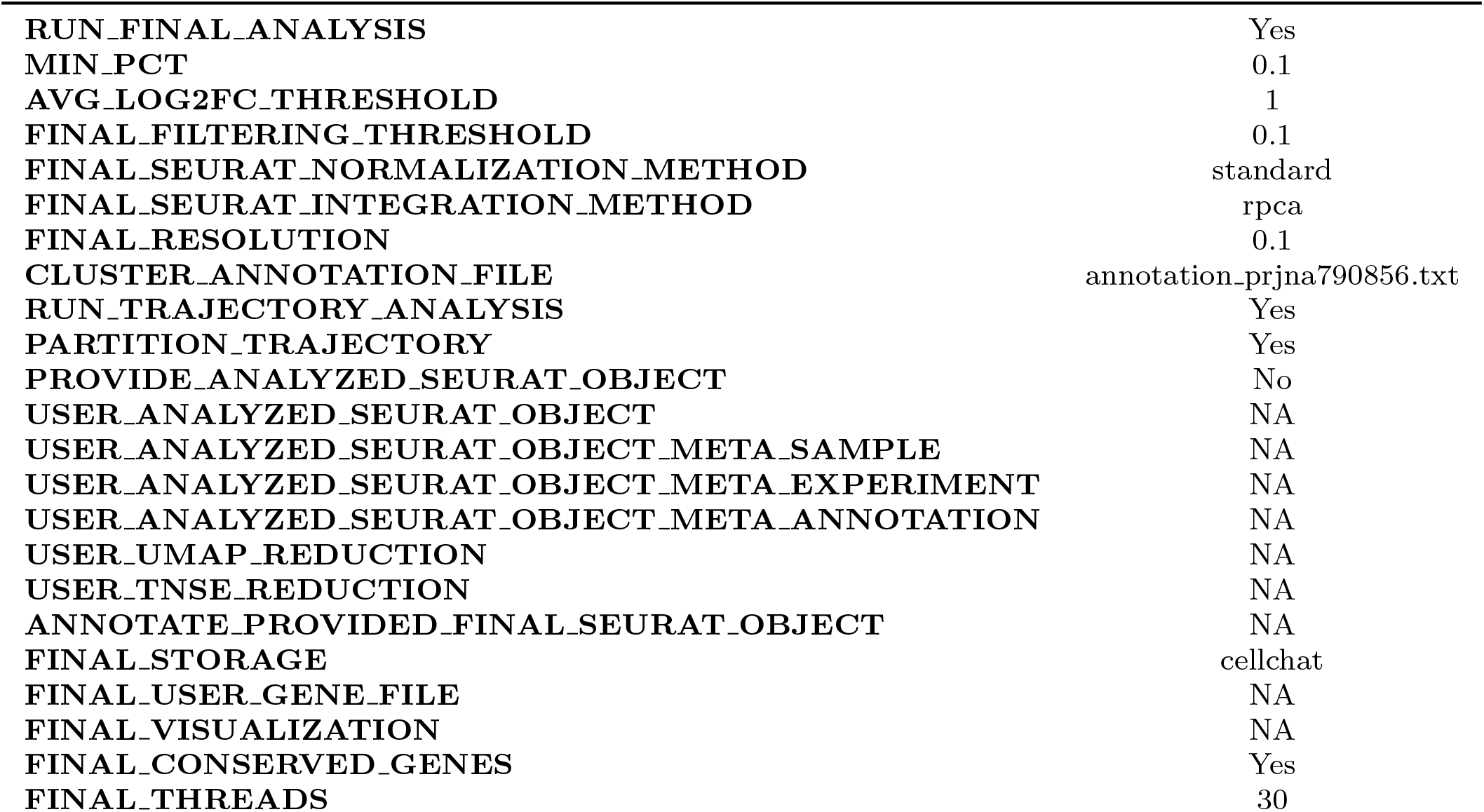
YAML Parameters used in the analysis of PRJNA790856 (GEO: GSE191288) for post-annotation analysis.

**Table 6.**
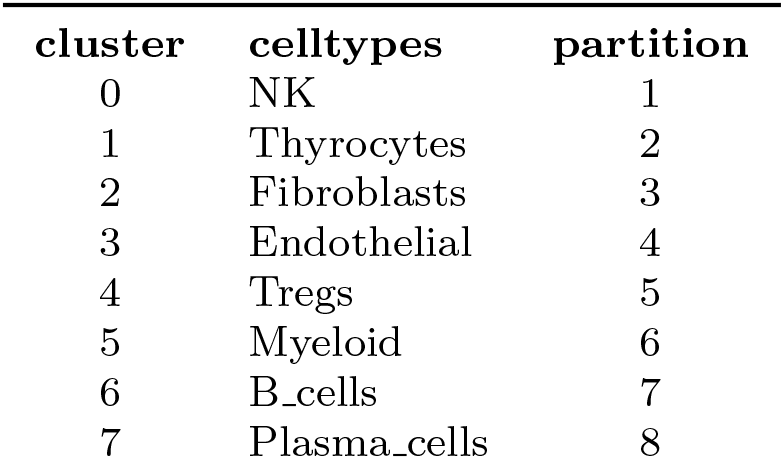
Example annotation file.

DEGs for each annotated cell type were found by comparing transcriptional differences across experimental conditions. Within each cluster, each gene was evaluated provided it was expressed in at least 10% of cells in both experimental groups (`MIN.PCT: 0.1`). The resulting DEGs for each cluster were filtered to retain only those with an average log2 fold change of 1 or higher (`AVG\_LOG2FC\_THRESHOLD`, and GSEA was run on the DEGs for each cluster.

A CellChat object (`FINAL\_STORAGE: cellchat`) was created to demonstrate the potential for downstream analysis of cell-cell communication and to include the computational time for benchmarking.

#### 5.2.1 Final analysis report

An overview of the post-annotation analysis report (generated for the GSE191288 dataset) is given in Figure 6. The SWANS post-annotation report displays the sample metadata and a list of which YAML parameters were used for the analysis (Figure 6A.), a UMAP with annotated clusters (Figure 6B.), an annotated UMAP split by experimental condition, an annotated bar-plot of cell proportions in each cluster for both experimental conditions, and an accompanying table that shows the number of cells (and their proportions) in each cluster across both experimental conditions (Figure 6C.). The report also includes tables of DEGs for between experimental conditions in each annotated cluster, and two tables of GSEA results (one for each of the ranking metrics: average log2 fold change and adjusted p-value) (Figure 6D.). While the report includes all GSEA results, associated images are not included due to the unknown variability in the number of results returned. These images can be found in the ‘pathways/figures’ directory within the output files. An example plot can be found in Figure 7. Additionally, there are three z-score tables (by cluster, by cluster split by experiments, and by cluster split by experiment and sample) (Figure 6E.). An annotated UMAP and barplot of cell proportions split by sample are also displayed in the report, along with a table of the number cells and their proportions (Figure 6F.). The report also includes a heatmap of the top 75 most variable genes and a table with a list of the 3000 most variable genes (Figure 6G.), a table of the conserved genes in each cluster (Figure 6H.), a UMAP colored by cell cycling phase Figure 6I., and lastly, a UMAP of the Monocle3 trajectory analysis (Figure 6J.). Trajectory analysis is most suitable for cell types that undergo dynamic transitions, such as stem and progenitor cells, immune cells during activation, cancer cells undergoing metastasis, or cells in developmental or differentiation processes, where continuous changes in gene expression or cell states can be tracked over time. While our pipeline includes the option to generate a Monocle3 object, we note that this method may not be suitable for all datasets depending on cell type composition and biological context. A full version of the HTML report in PDF format is located in Supplementary materials and includes expanded table descriptions (Supplementary 3).

**Fig. 6.**
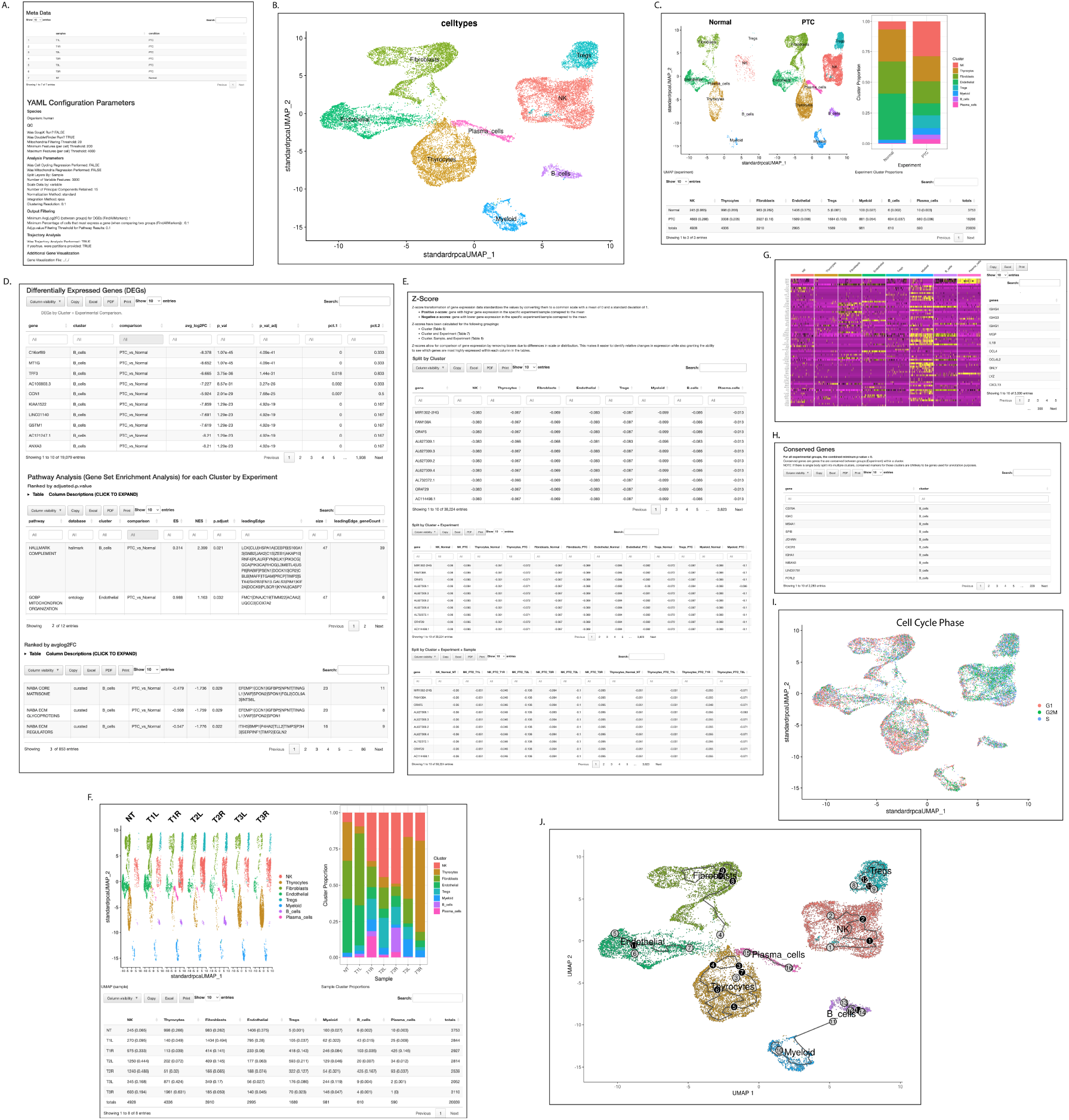
Overview of final report. A. Sample name and experiment assignment (top) and overview of YAML user-assigned parameters used during analysis (bottom). B. Annotated UMAP. C. Annotated UMAP split by experimental condition (left), annotated barplot of cell proportions by cluster and experimental condition (right), and table depicting the number and proportion of cells in each cluster and split by experimental condition (bottom). D. Table of within cluster (by experiment) DEGs (top), filtered GSEA results for each cluster when DEGs were ranked by adjusted p value (middle), and filtered GSEA results for each cluster when DEGs were ranked by average log2 fold change (bottom). E. Gene transcript z-score transformations by cluster (top), cluster and experiment (middle), and cluster, experiment (full table not shown), and sample (full table not shown) (bottom). F. Annotated UMAP split by sample (left), annotated barplot of cell proportions by cluster and sample (right), and a table containing the number and proportion of cells in each cluster when split by sample (bottom). G. Heatmap of the 75 most variable genes (left) and gene names of the 3000 most variable genes (left). H. Table of conserved genes by cluster. I. UMAP colored by cycling phase. J. Figure displaying the trajectory analysis graph from Monocle3 over a UMAP of the annotated clusters. The full report is interactive and in HTML format; an example of this report in PDF format can be found in Supplementary 3.

**Fig. 7.**
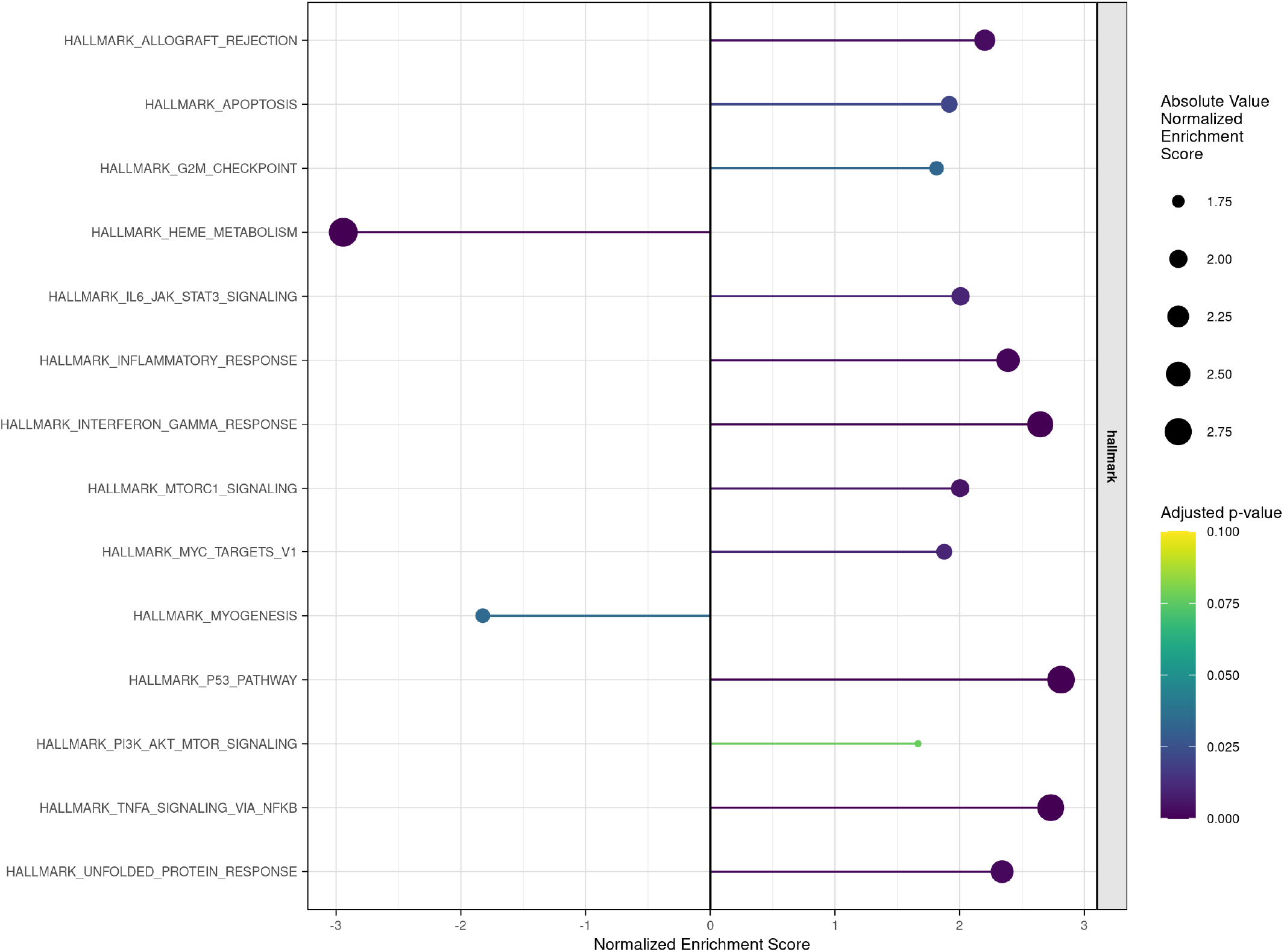
GSEA plot of B cell cluster results comparing Papillary Thyroid Carcinoma (PTC) cells and Normal cells, with DEGs ordered by average log2 fold change, using the Hallmark Molecular Signatures Database (MSigDB).

#### 5.2.2 Benchmarking

Snakemake’s benchmarking feature allows users to track and record the resource usage (such as time and memory consumption) of each step in a workflow. A benchmark report was generated for PRJNA790856, showing the computational cost of each individual rule in the SWANS workflow (Figure 8). A full version of the report HTML report in PDF format is located in Supplementary materials (Supplementary 4).

**Fig. 8.**
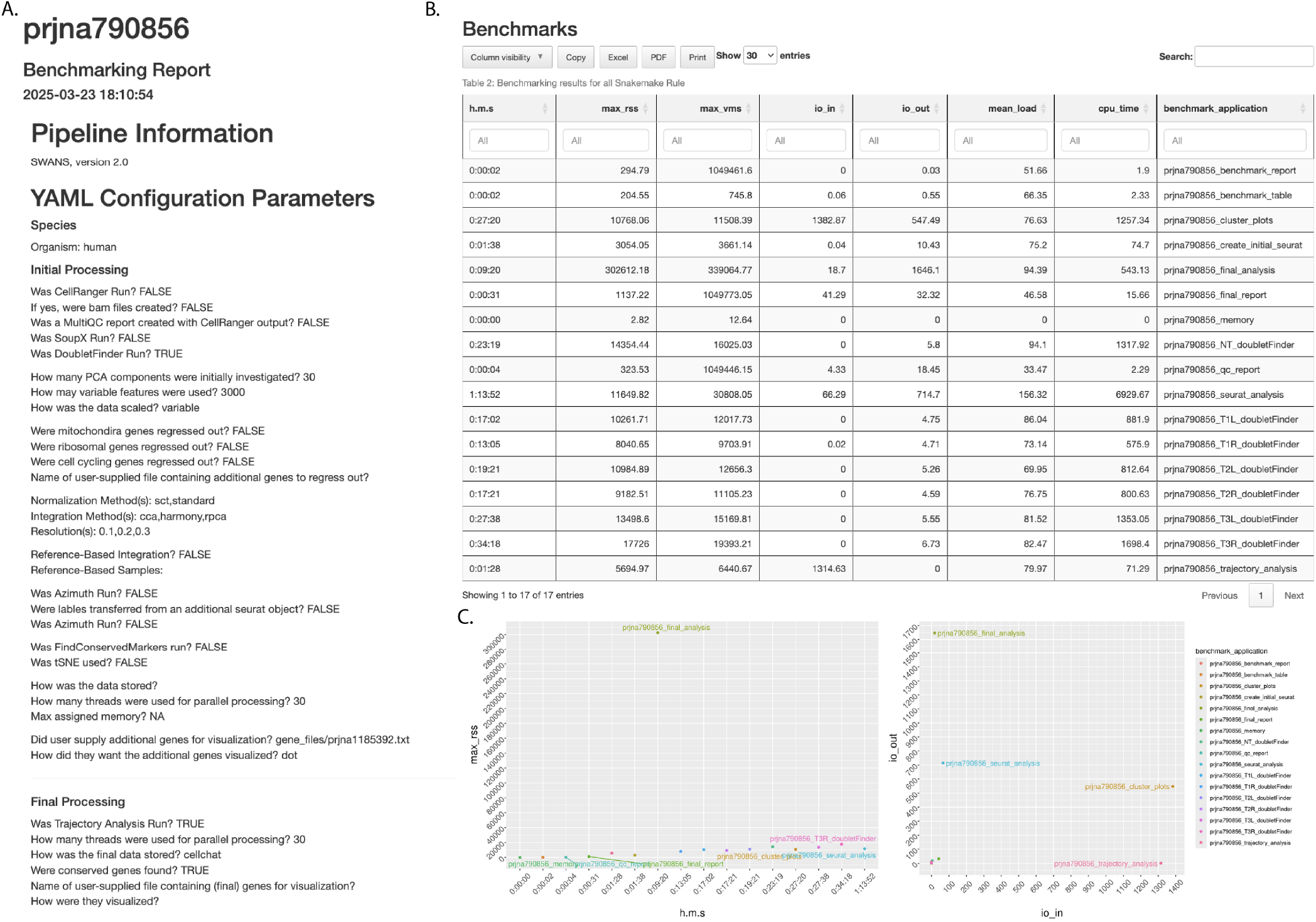
Benchmarking report. A. Overview of parameters used in analysis. B. Resources used for each rule in SWANS pipeline. C. Plots of run time vs max memory and number of MB read vs. number of MB written. Performance metrics: ‘s’ (running time in seconds), ‘h:m:s’ (running time in hour, minutes, seconds format), ‘max_rss’ (maximum Resident Set Size, the non-swapped physical memory used), ‘max_vms’ (maximum Virtual Memory Size, total virtual memory used), ‘max_uss’ (Unique Set Size, memory unique to a process), ‘max-pss’ (Proportional Set Size, memory shared with other processes, divided evenly, Linux only), ‘io_in’ (cumulative megabytes (MB) read), ‘io_out’ (cumulative MB written), ‘mean_load’ (CPU usage over time, divided by total running time), and ‘cpu_time’ (summed CPU time for user and system). The full report is in HTML format; an example of this report in PDF format can be found in Supplementary 4.

To thoroughly benchmark the pipeline, we downloaded eight publicly available datasets, which were then processed through the SWANS pipeline to evaluate its performance across a variety of sample sizes, tissues, tissue states, and starting data format (Figure 9). The pipeline was executed on a Linux operating system (Red Hat Enterprise Linux 8.10 (OOTPA)), ensuring a robust computational environment for the analysis. To provide complete transparency and reproducibility, all the specific parameters used to process these datasets are detailed in Table 7. This benchmarking approach not only tests the e”ciency of SWANS but also demonstrates its versatility when applied to diverse datasets.

**Table 7.**
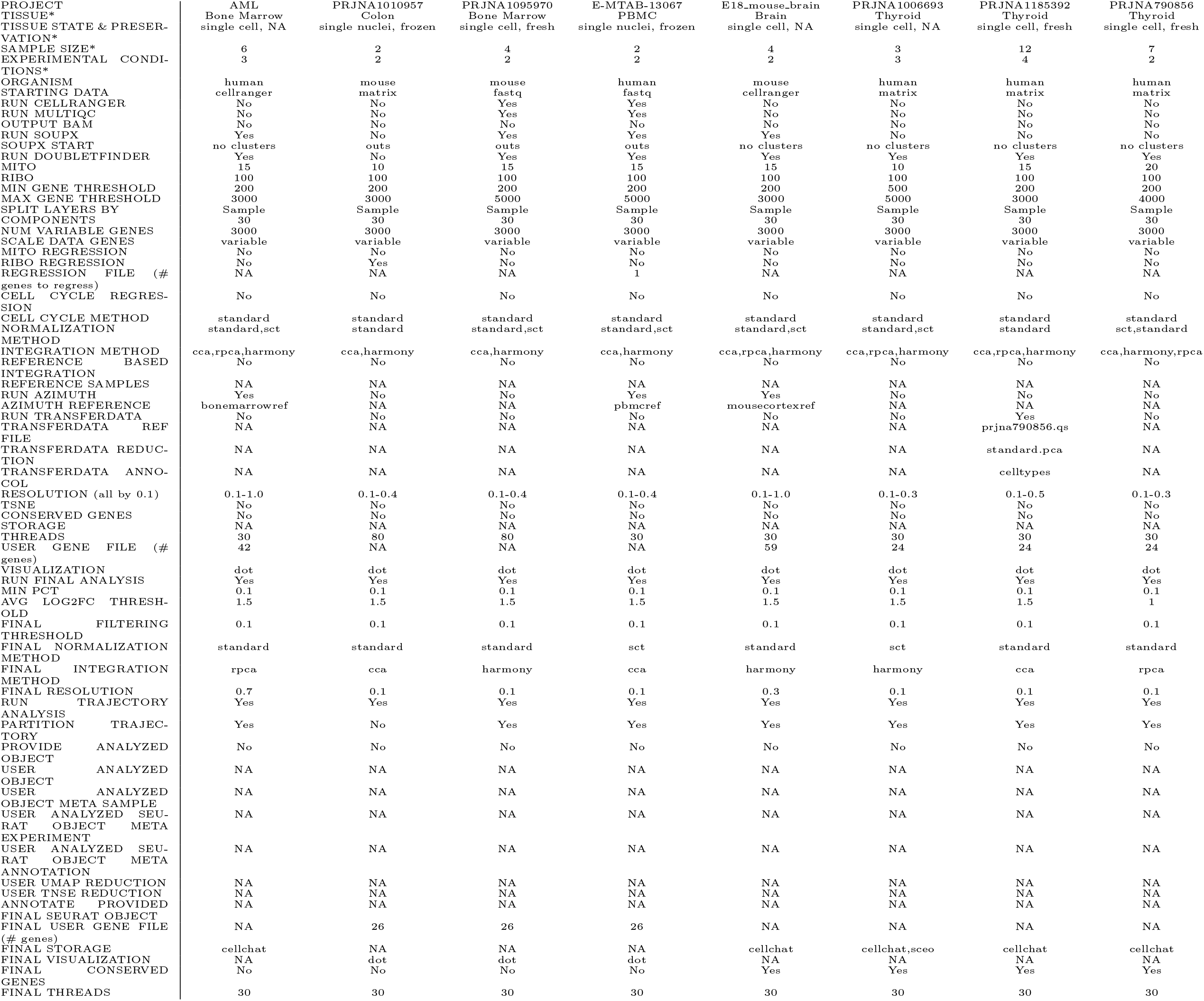
Overview of all YAML configurations used to process each publicly available dataset. *Added to table for comparative purposes. Some words in leading column have been shorted or removed for fitting purposes.

**Fig. 9.**
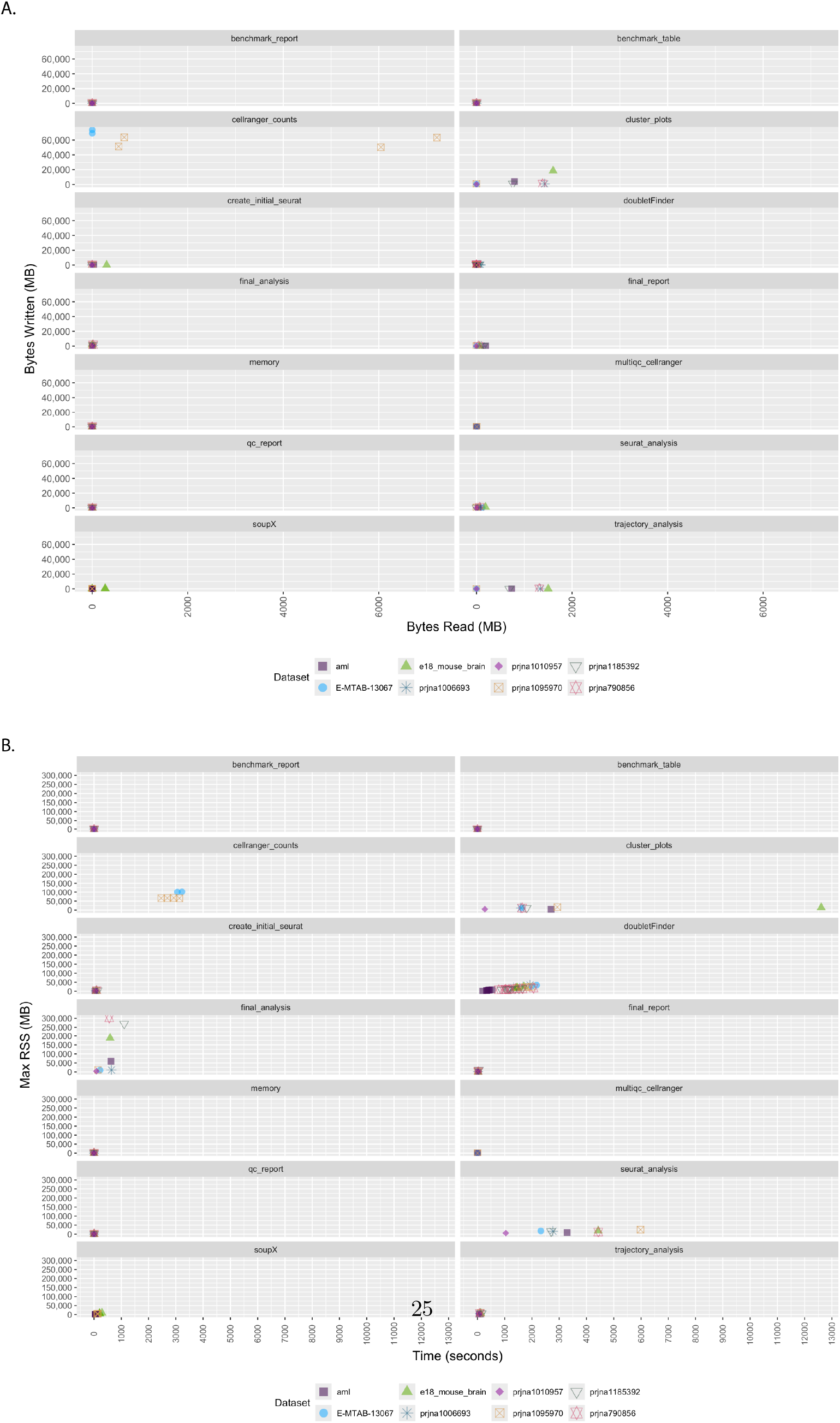
Resource and time measurements for each step in the preliminary and final analysis for eight publicly available datasets. A. Megabytes (MB) read in vs MB written for each rule and dataset. B. Time in seconds used for each rule and dataset.

## 6 Conclusion

SWANS is a customizable pipeline for analysis of scRNA-seq/snRNA-seq data that allows the user to compare multiple analysis schemas in an interactive report. SWANS uses Snakemake for workflow management, which simplifies and automates complex workflows by providing a clear, reproducible pipeline structure, e”cient parallelization, scalability, and compatibility with various computational environments. SWANS can be customized (via configuration files) to tailor the analysis to specific experimental conditions and biological question(s).

A key motivation behind SWANS was to enhance communication between bioin-formaticians and their collaborators. With SWANS, bioinformaticians can easily share complex data and results with investigators to promote meaningful discussions on how to proceed with downstream analysis. In the interest of accessibility and collaboration, we implement open source software tools and packages which are provided in Docker image, and we provide our source code for SWANS in GitHub repository.

## Supporting information

Supplementary pdfs 1 - 4

## Supplementary information

## Acknowledgments

We gratefully acknowledge the entire Franco Laboratory for their thoughtful discussions and constructive feedback throughout the development of this tool and the subsequent reports. We also thank the Heuckeroth Lab for their invaluable feedback and support during the development process. A very special note of appreciation to Eric Howden for his continual and gentle presence during the coding development.

## Declarations

- Ethics approval and consent to participate: Not applicable
- Consent for publication: Not applicable
- Availability of data and materials: The datasets analyzed during the current study are available here: 10X Genomics [49–59], ArrayExpress dataset (E-MTAB-13067) [48], and NCBI GEO datasets [60–64].
- Conflict of interest/Competing interests: The authors declare that they have no competing interests.
- Funding: Authors received support for this work from the Department of Biomedical and Health Informatics and the CHOP Research Institute at The Children’s Hospital of Philadelphia. This work was supported in part by Department of Defense award W81XWH2210655 (awarded to ATF). The funders had no role in study design, data collection and analysis, decision to publish, or preparation of the manuscript.
- Author contribution: KB: Coding, testing, software instructions, co-wrote manuscript; EW: Coding, compiling Docker image, reviewed manuscript; SJP: code testing, wrote/reviewed manuscript; MG: Coded initial Shiny app; ROH: Supplied dataset, reviewed manuscript; JCRF & ATF: Provided feedback on visualization/reports, reviewed manuscript; ERR conceived of idea and wrote all original code on which this version is based, testing, software instructions, co-wrote manuscript. All authors read and approved the final manuscript.
- Materials availability: Not applicable
- Code availability: Source code and documentation are available at https://github.com/FrancoResearchLab/SWANS. A Docker image for running SWANS is available at https://hub.docker.com/r/francothyroidlab/swans.

