## Supplementary pdfs 1 - 4 for "SWANS: A highly configurable analysis pipeline for single-cell and single-nucleus RNA-sequencing data"

**Supplementary information.**

### 1 Supplementary PDF 1: QC Report

### prjna790856

#### Quality Control Report

2025-03-23 13:52:19

### Pipeline Information

SWANS, version 2.0

### YAML Configuration Parameters

##### Species

Organism: human

### QC

Was SoupX Run? FALSE
Was DoubletFinder Run? TRUE
Mitochondria Filtering Threshold: 20
Ribosomal Filtering Threshold: 100
Minimum Features (per cell) Threshold: 200
Maximum Features (per cell) Threshold: 4000

---

### DoubletFinder

#### Sample: T1L

Total number of cells: 4042
Doublet rate: 0.031
Total number of doublets: 118
Number of cells after doublet removal: 3924

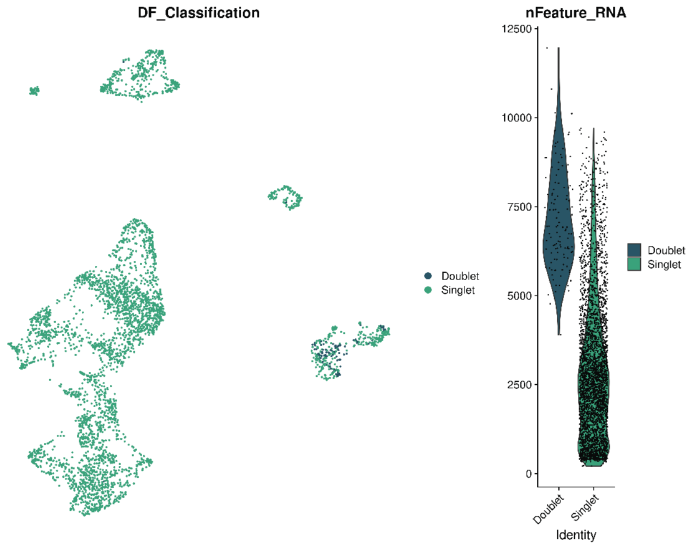

### Sample: T1R

Total number of cells: 4155

Doublet rate: 0.031

Total number of doublets: 113

Number of cells after doublet removal: 4042

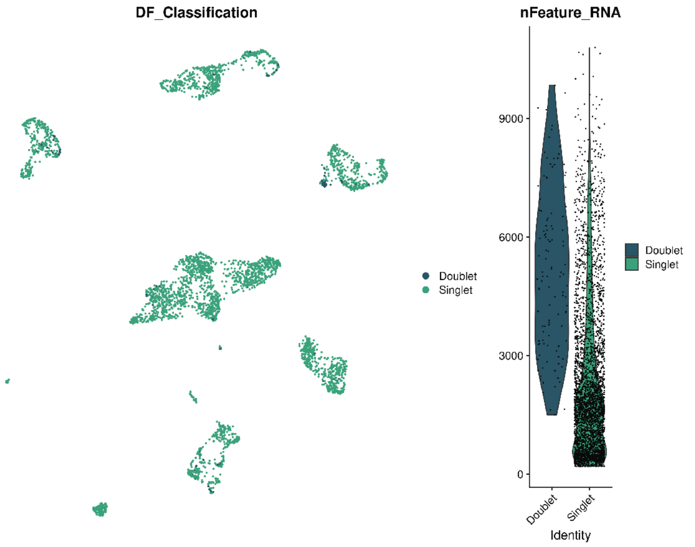

---

### Sample: T2L

Total number of cells: 4390

Doublet rate: 0.031

Total number of doublets: 131

Number of cells after doublet removal: 4259

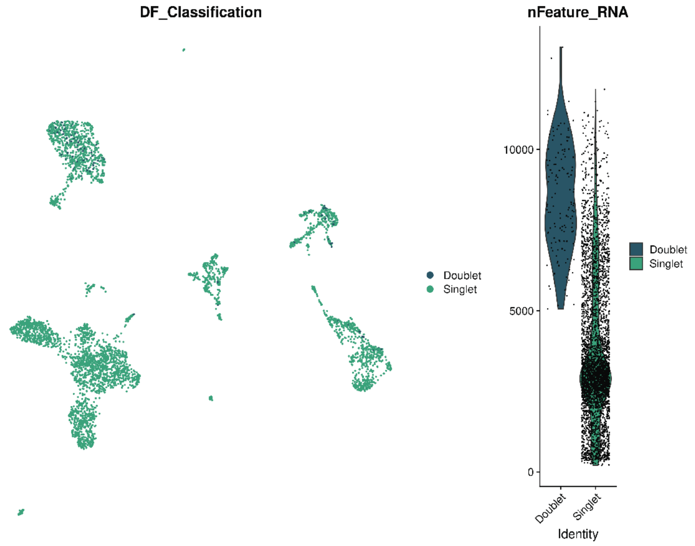

### Sample: T2R

Total number of cells: 3543

Doublet rate: 0.031

Total number of doublets: 104

Number of cells after doublet removal: 3439

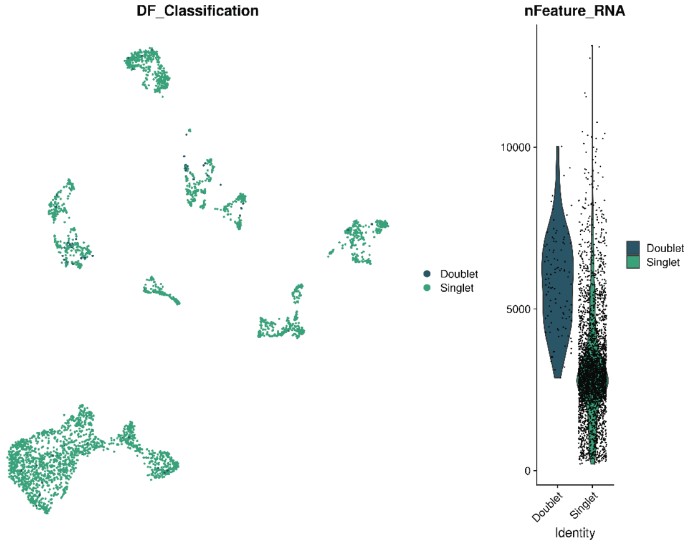

---

### Sample: T3L

Total number of cells: 4901

Doublet rate: 0.039

Total number of doublets: 188

Number of cells after doublet removal: 4713

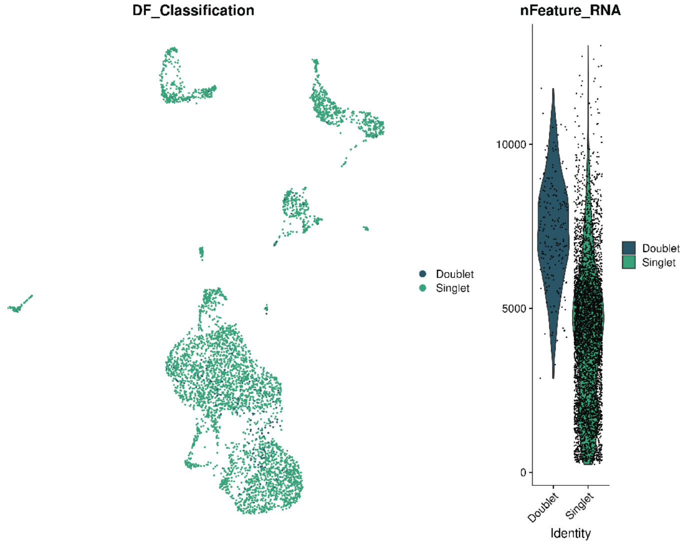

### Sample: T3R

Total number of cells: 6364

Doublet rate: 0.046

Total number of doublets: 291

Number of cells after doublet removal: 6073

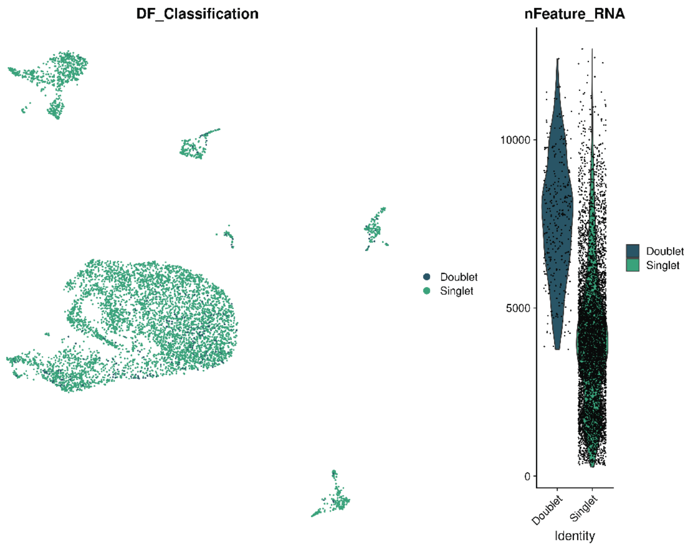

---

### Sample: NT

Total number of cells: 5859

Doublet rate: 0.046

Total number of doublets: 227

Number of cells after doublet removal: 5632

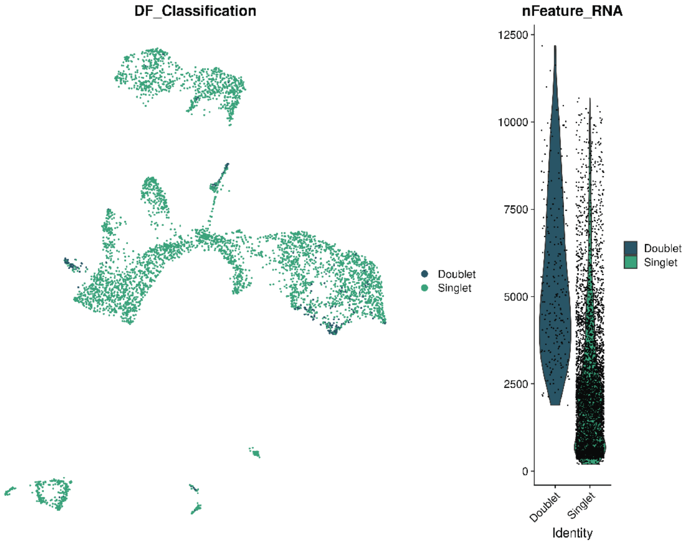

### Seurat QC plots

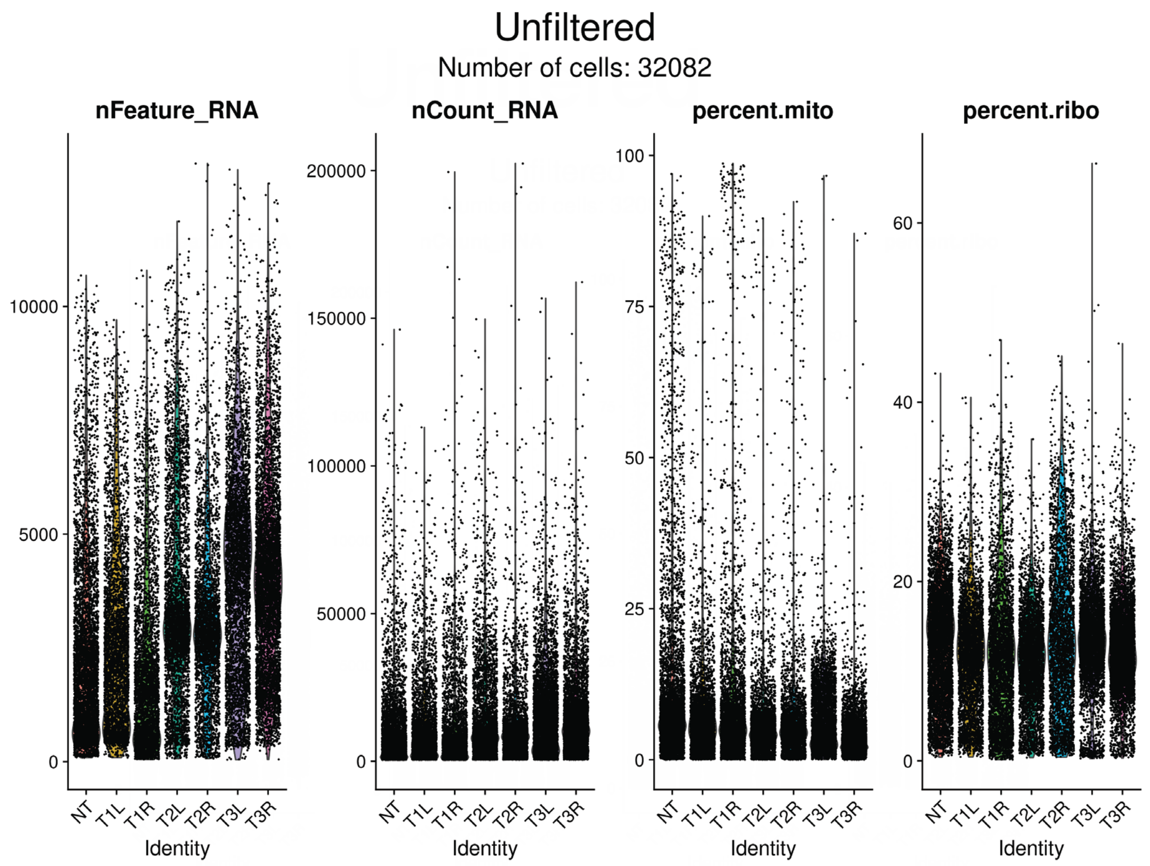

Data before filtering

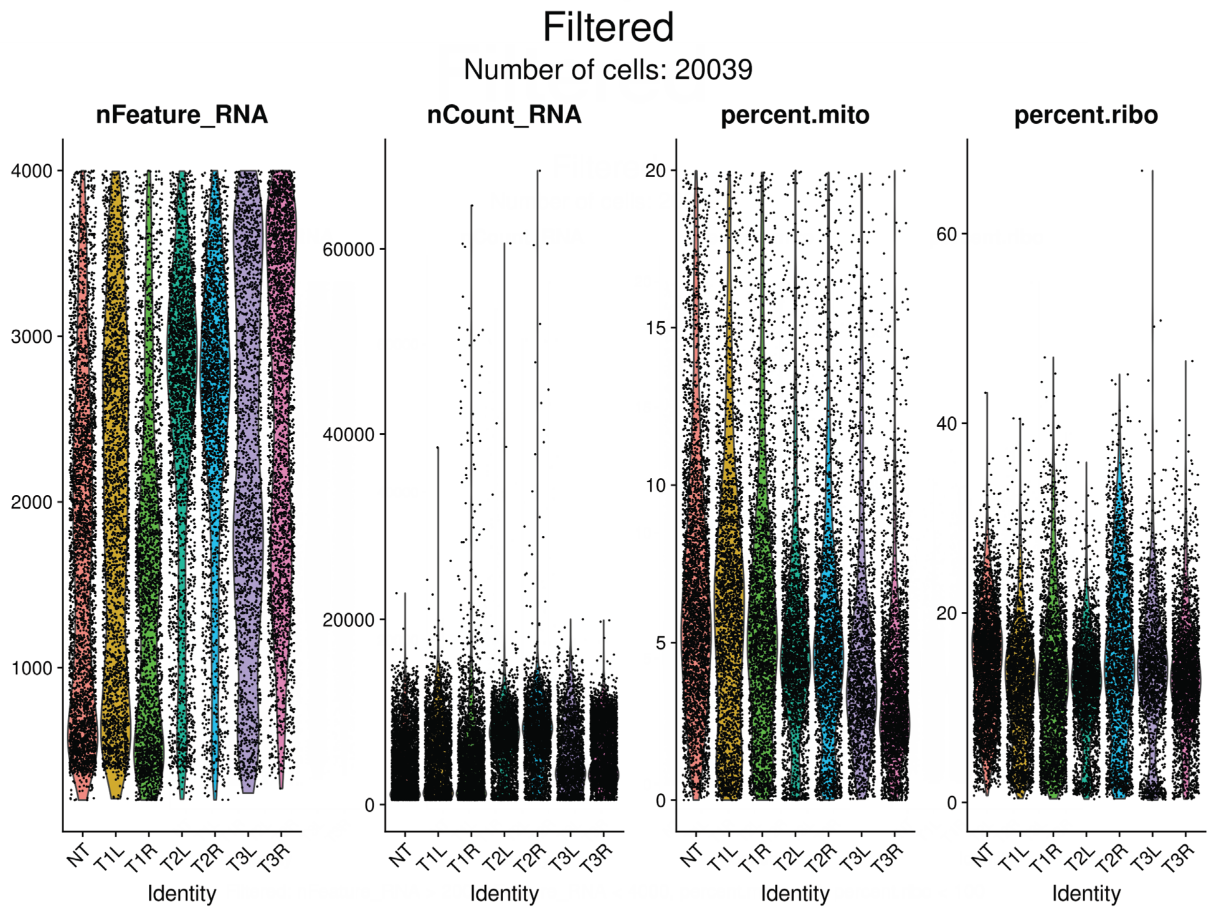

Filtered: nFeature_RNA > 200, nFeature_RNA < 4000, percent.mito < 20, percent.ribo < 100

### Seurat QC plots

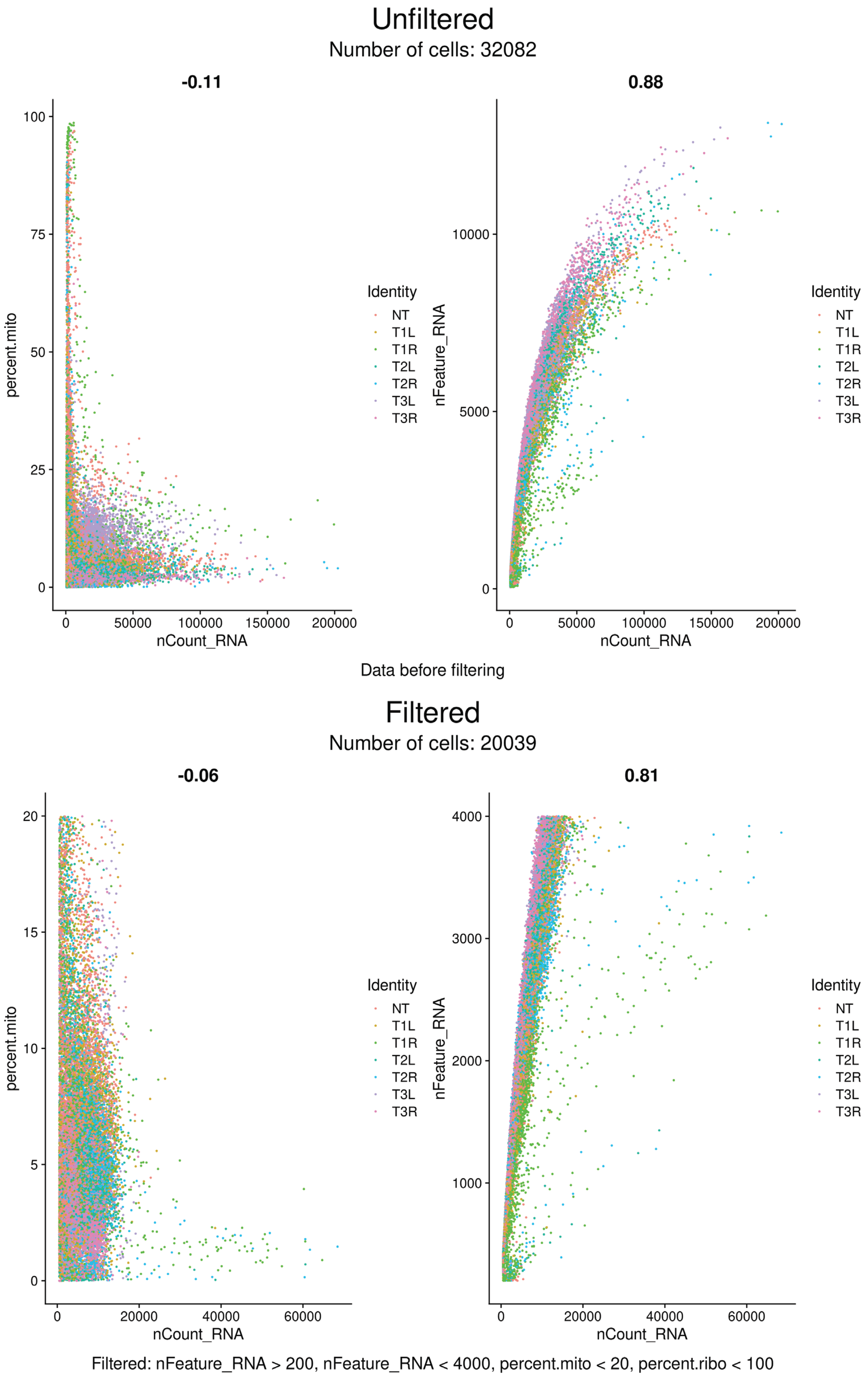

### Seurat QC plots

#### Unfiltered

Number of cells: 32082

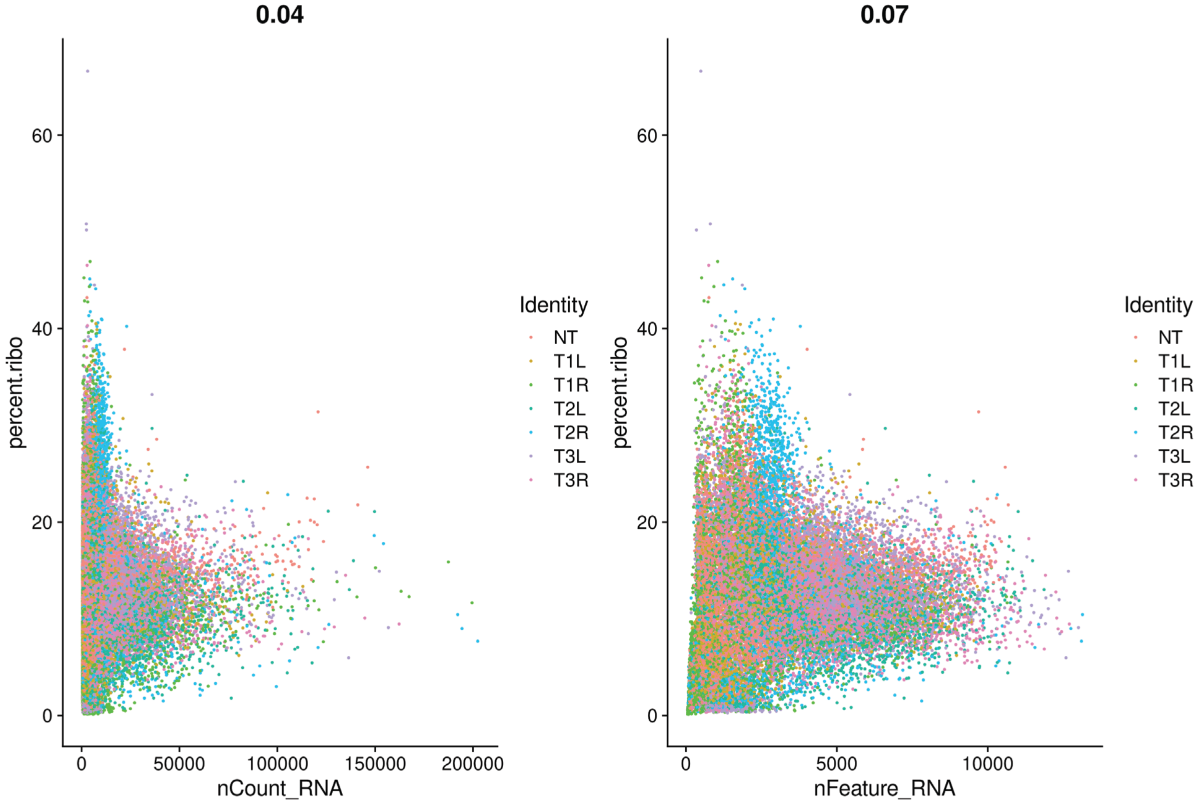

Data before filtering

#### Filtered

Number of cells: 20039

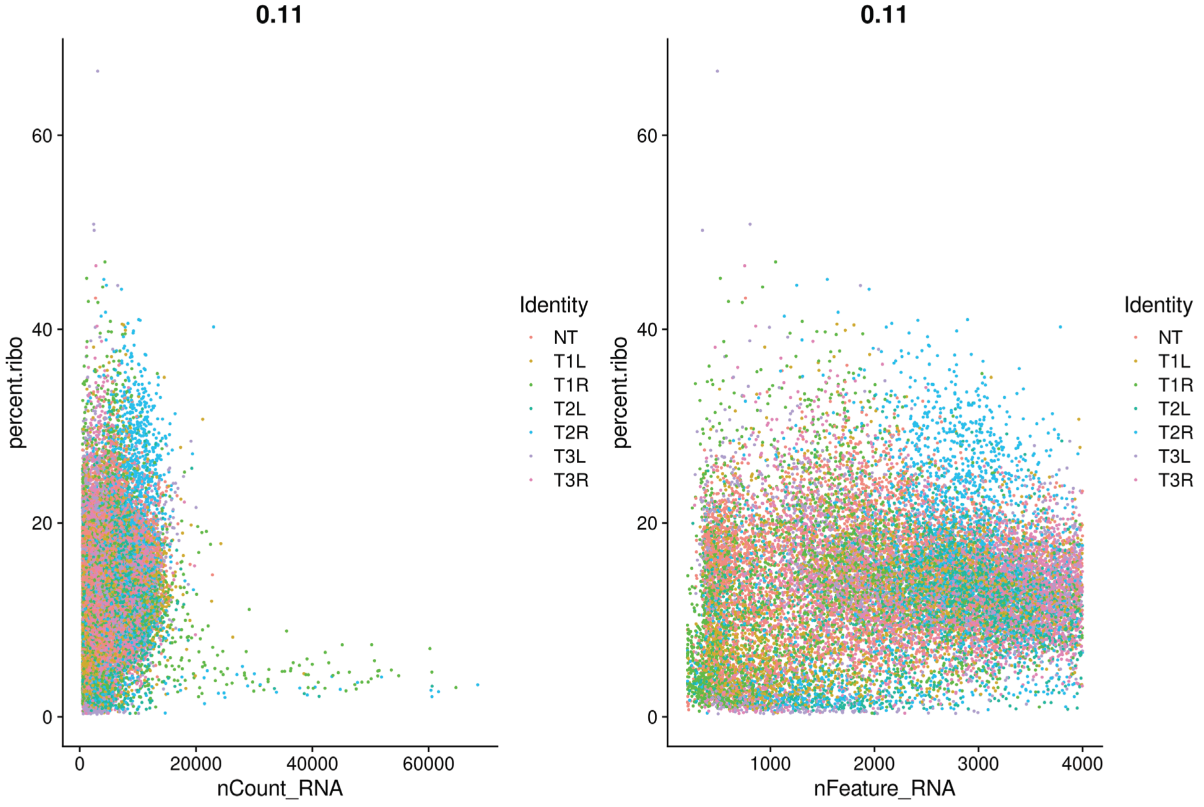

Filtered: nFeature_RNA > 200, nFeature_RNA < 4000, percent.mito < 20, percent.ribo < 100

#### 2 Supplementary PDF 2: Additional PDF Output

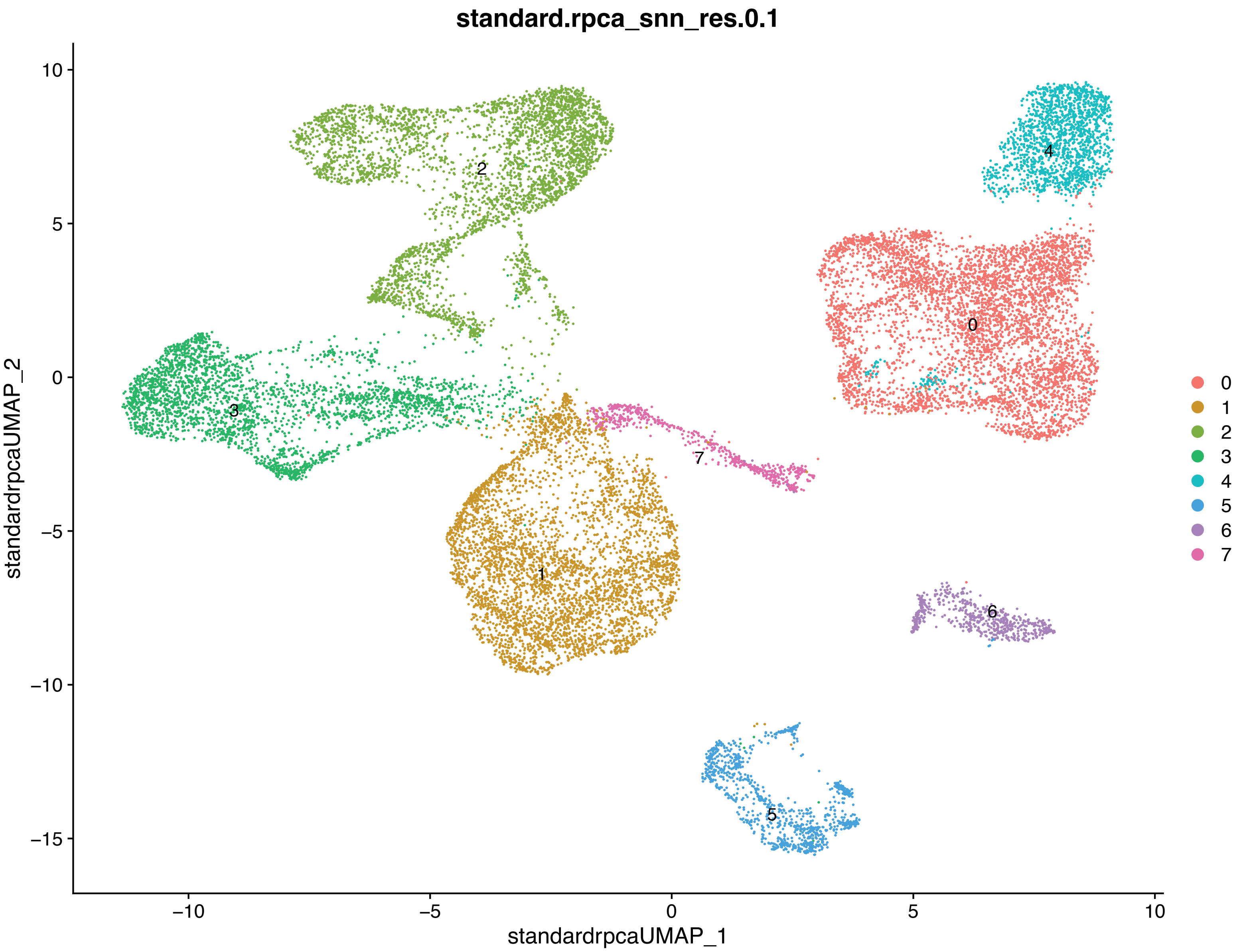

standard.rpca_snn_res.0.1
standardrpcaUMAP_2
standardrpcaUMAP_1
10
5
0
−5
−10
−15
−10
−5
0
5
10
0
1
2
3
4
5
6
7

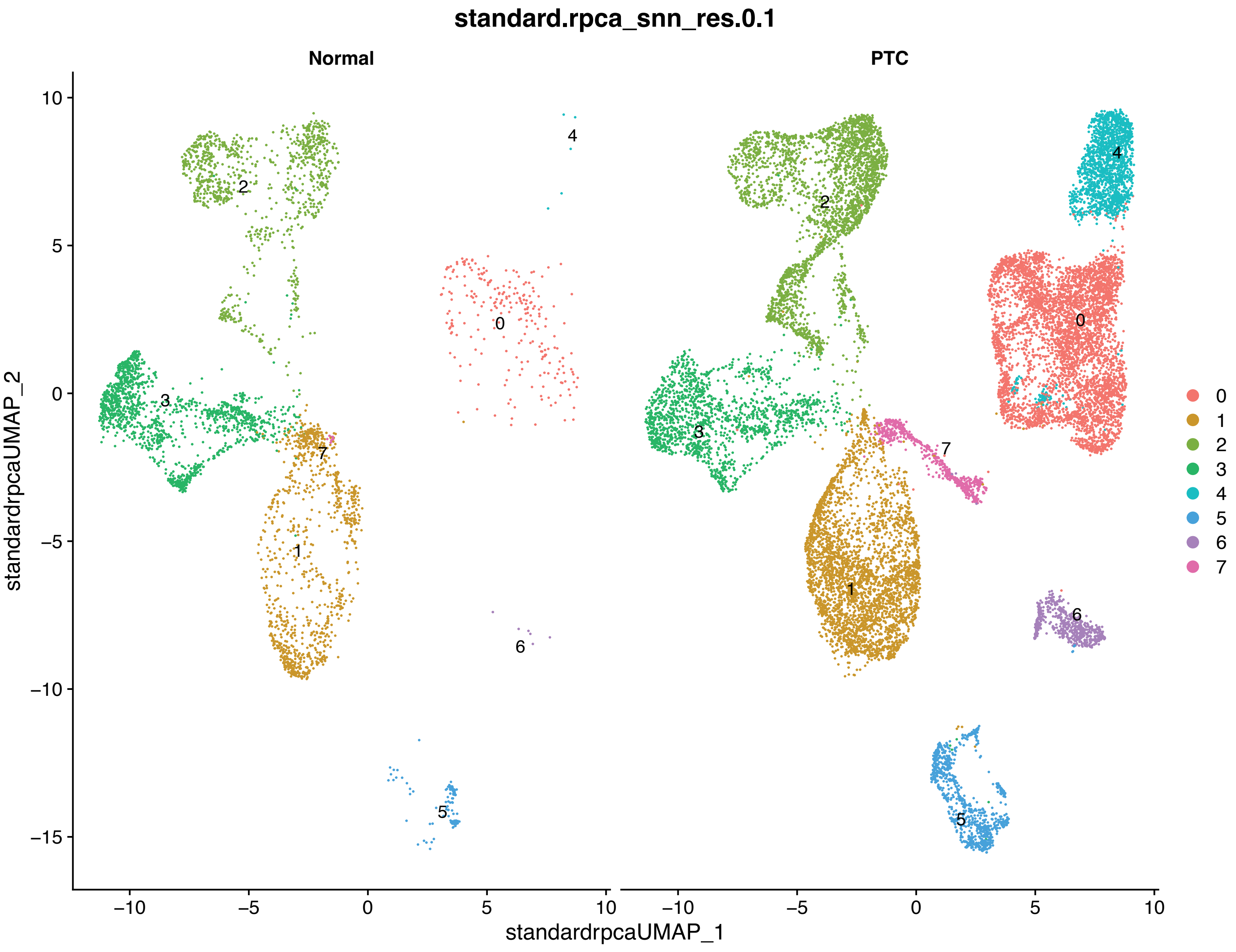

standard.rpca_snn_res.0.1
Normal
PTC
standardrpcaUMAP_2
standardrpcaUMAP_1
10
5
0
−5
−10
−15
−10
−5
0
5
10
−10
−5
0
5
10
0
1
2
3
4
5
6
7

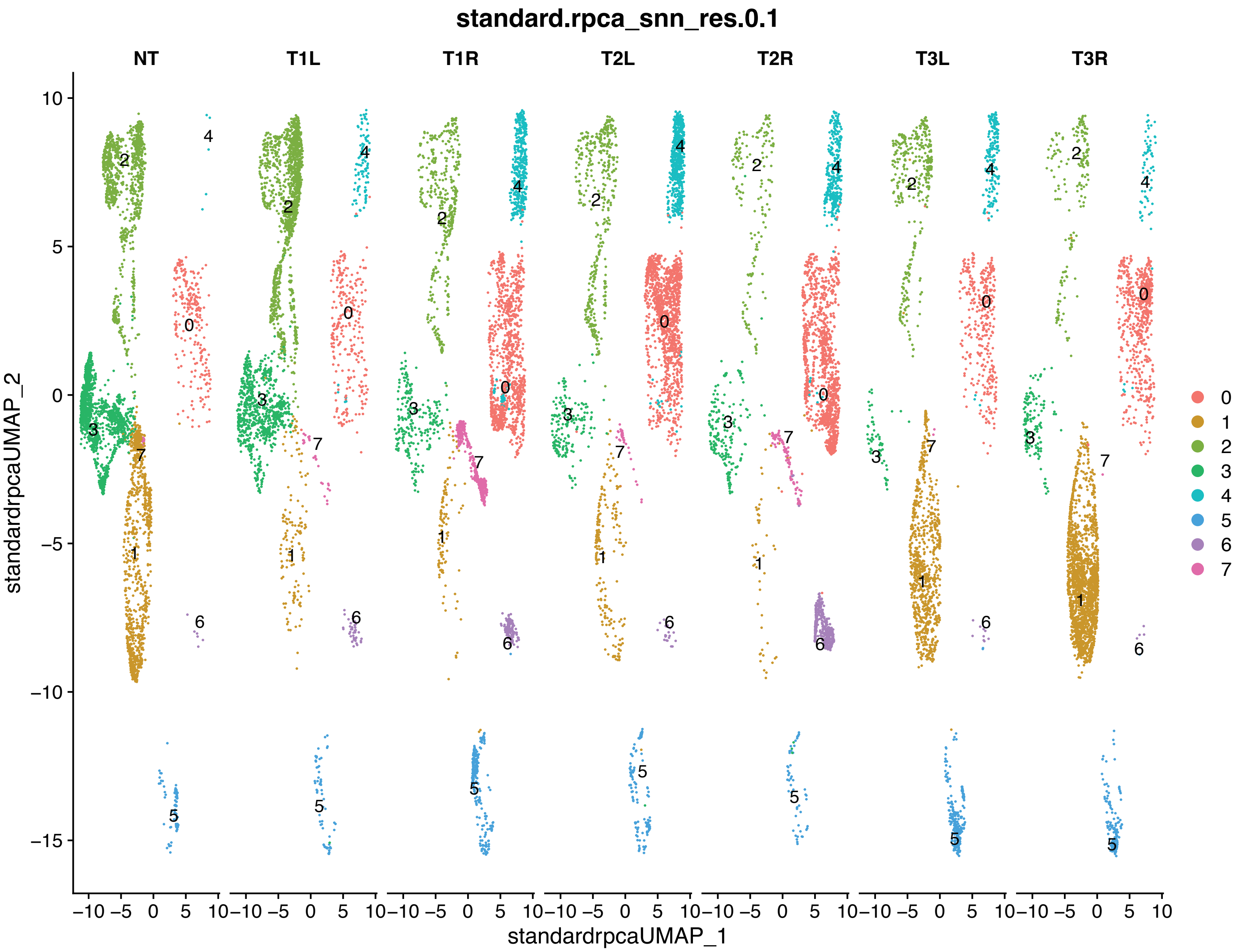

standard.rpca_snn_res.0.1
NT
T1L
T1R
T2L
T2R
T3L
T3R
standardrpcaUMAP_2
standardrpcaUMAP_1
10
5
0
−5
−10
−15
−10 −5 0 5 10
0
1
2
3
4
5
6
7

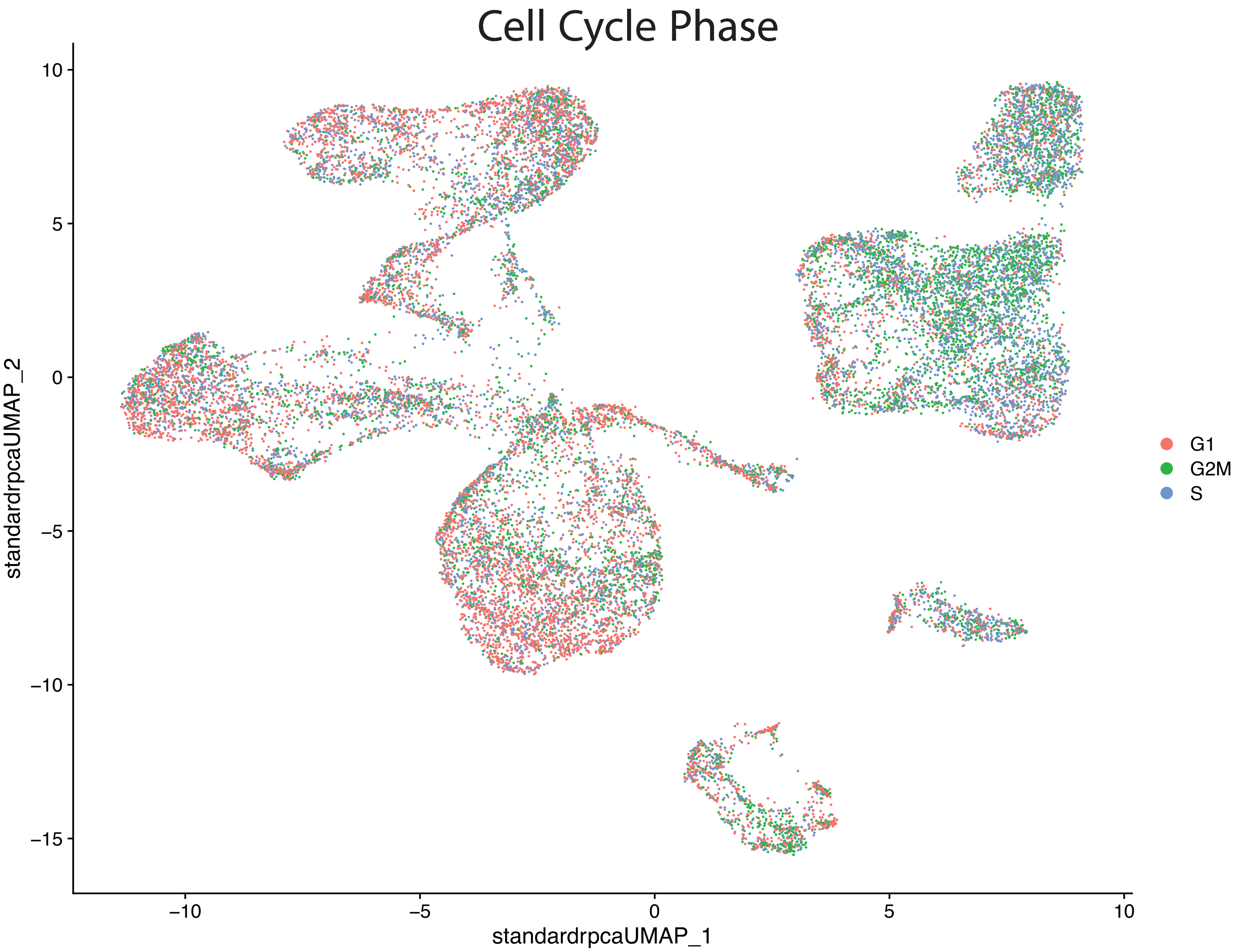
Cell Cycle Phase
10
5
0
−5
−10
−15
standardrpcaUMAP_2
−10
−5
0
5
10
standardrpcaUMAP_1
G1
G2M
S

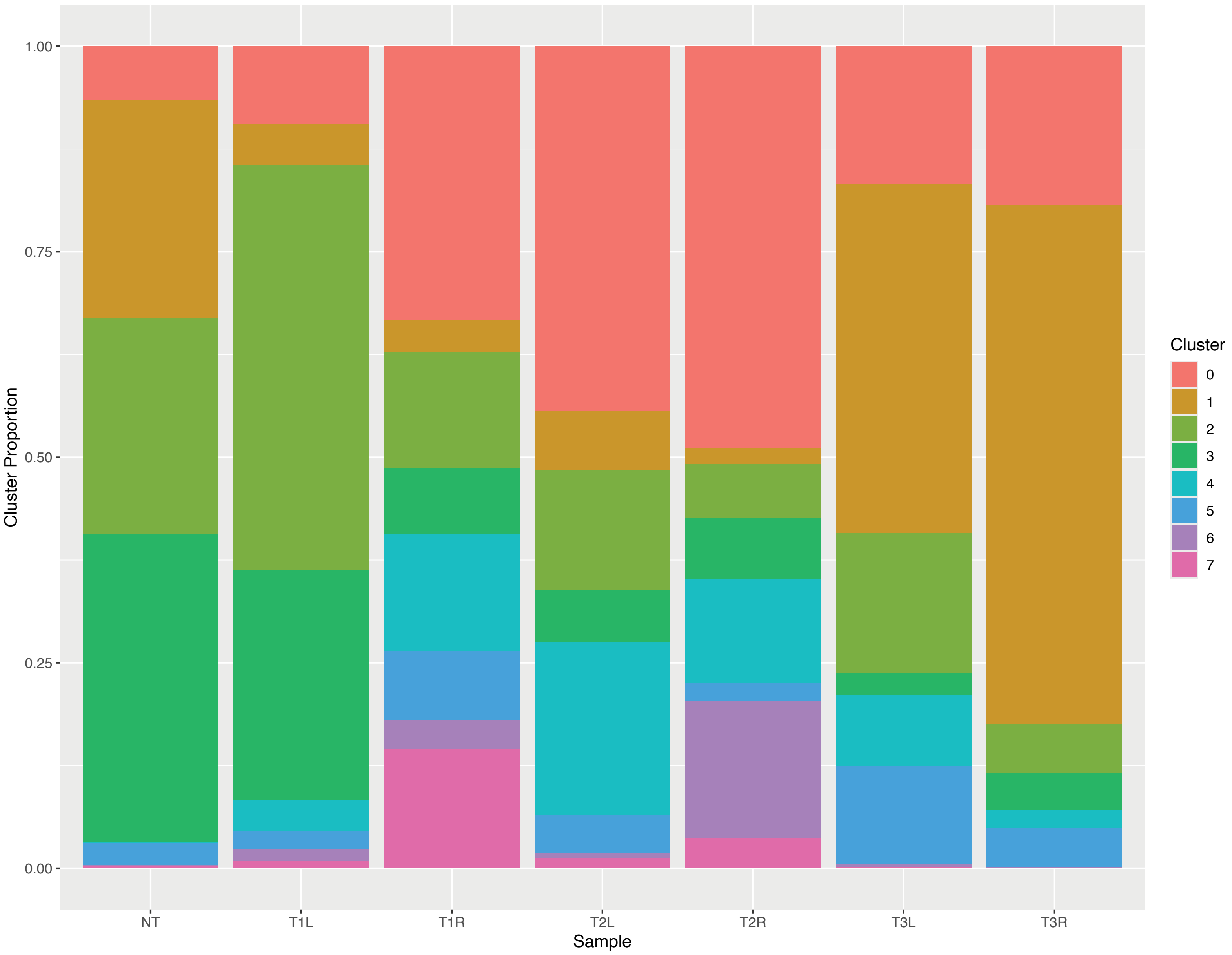

1.00
0.75
0.50
0.25
0.00
Cluster Proportion
NT
T1L
T1R
T2L
T2R
T3L
T3R
Sample
Cluster
0
1
2
3
4
5
6
7

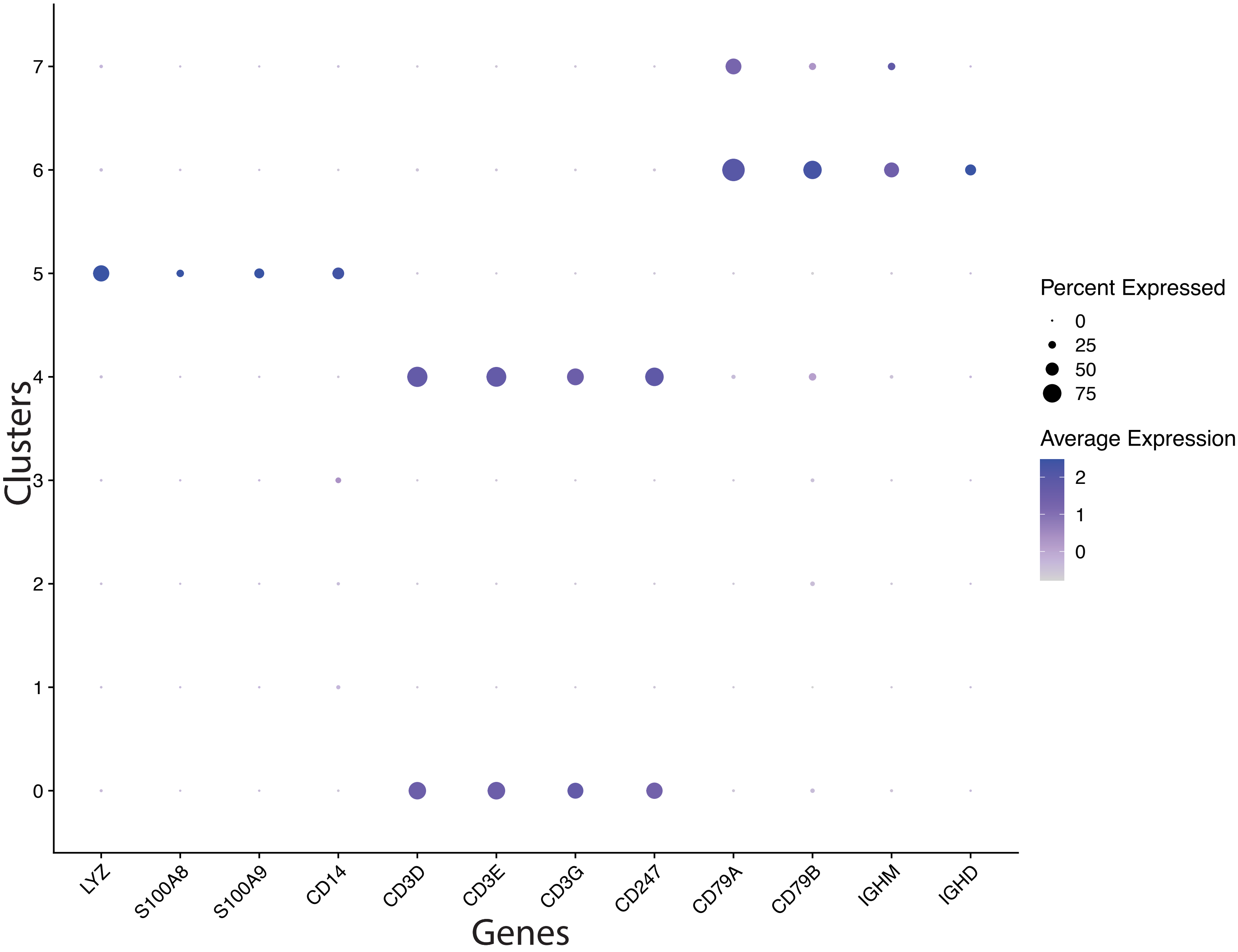

Clusters
7
6
5
4
3
2
1
0
LYZ
S100A8
S100A9
CD14
CD3D
CD3E
CD3G
CD247
CD79A
CD79B
IGHM
IGHD
Genes
Percent Expressed
0
25
50
75
Average Expression
2
1
0

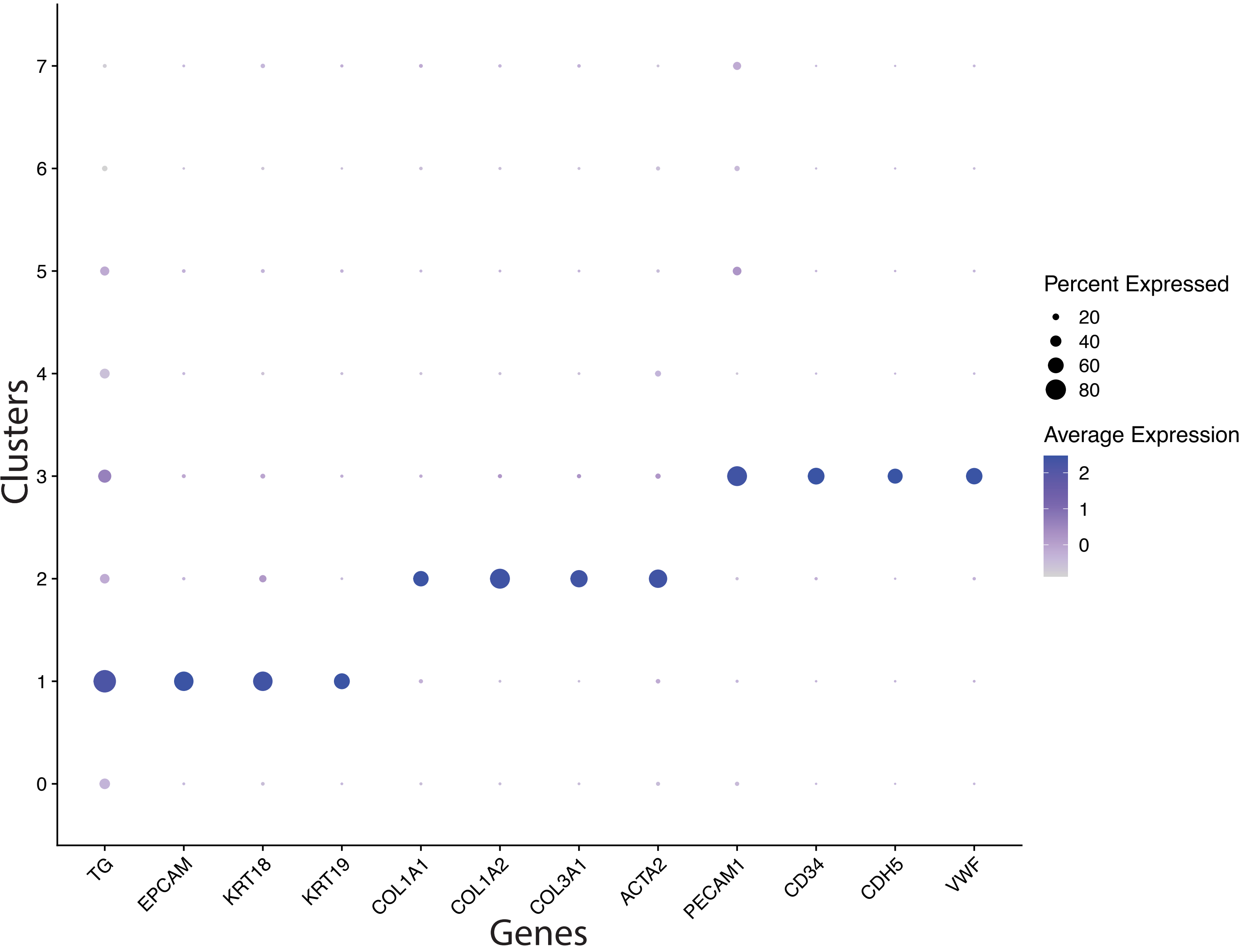

Clusters
7
6
5
4
3
2
1
0
TG
EPCAM
KRT18
KRT19
COL1A1
COL1A2
COL3A1
ACTA2
PECAM1
CD34
CDH5
VWF
Genes
Percent Expressed
20
40
60
80
Average Expression
2
1
0

#### 3 Supplementary PDF 3: Final Report

### prjna790856

E. Reichenberger

2025-04-14 16:45:59

#### SWANS

The cells were analyzed using SWANS, an automated and highly customizable single cell/nucleus RNA-seq analysis pipeline that relies heavily on Seurat, is managed by Snakemake, and must start with cellranger data.

#### Metadata

Show 10 entries Search:

Table 1: Sample Meta Data.

| | samples | condition |
|---|---|---|
| 1 | T1L | PTC |
| 2 | T1R | PTC |
| 3 | T2L | PTC |
| 4 | T2R | PTC |
| 5 | T3L | PTC |
| 6 | T3R | PTC |
| 7 | NT | Normal |

Showing 1 to 7 of 7 entries

Previous 1 Next

### YAML Configuration Parameters

##### Species

Organism: human

### QC

Was SoupX Run? FALSE
Was DoubletFinder Run? TRUE
Mitochondria Filtering Threshold: 20
Minimum Features (per cell) Threshold: 200
Maximum Features (per cell) Threshold: 4000

##### Analysis Parameters

Was Cell Cycling Regression Performed: FALSE
Was Mitochondria Regression Performed: FALSE
Split Layers By: Sample
Number of Variable Features: 3000
Scale Data by: variable
Number of Principal Components Retained: 15
Normalization Method: standard
Integration Method: rpca
Clustering Resolution: 0.1

##### Output Filtering

Minimum AvgLog2FC (between groups) for DGEs (FindAllMarkers): 1
Minimum Percentage of cells that must express a gene (when comparing two groups (FindAllMarkers)) : 0.1
Adj.p.value Filtering Threshold for Pathway Results: 0.1

##### Trajectory Analysis

Was Trajectory Analysis Performed: TRUE
If yes/true, were partitions provided: TRUE

##### Additional Gene Visualization

Gene Visualization File: ../../

### UMAP of Cluster Annotations

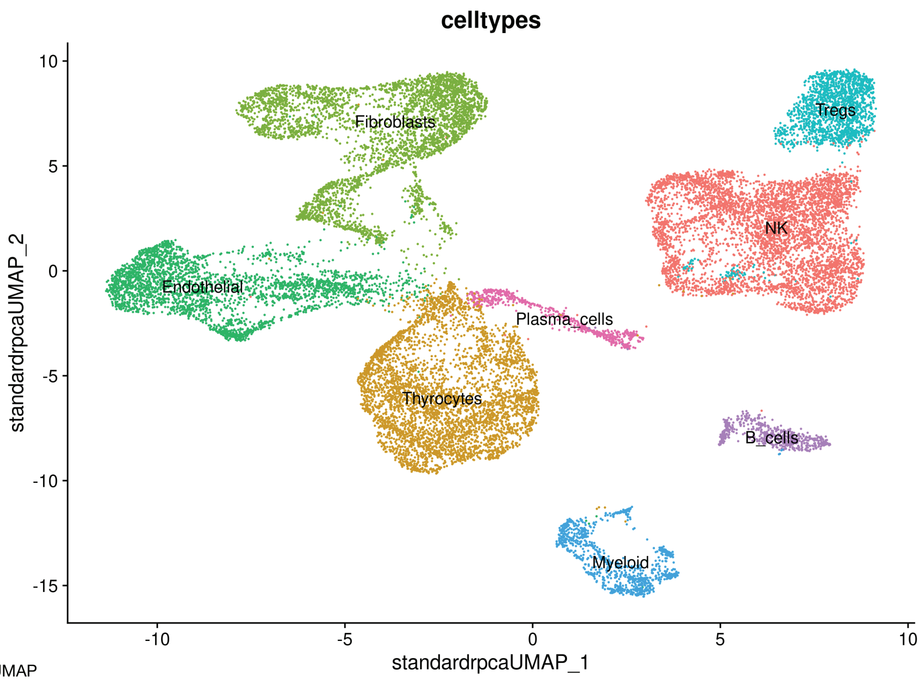

UMAP

### Experiment Comparisons

#### UMAP + Cell Counts (Proportions) for each Cluster by Experiment

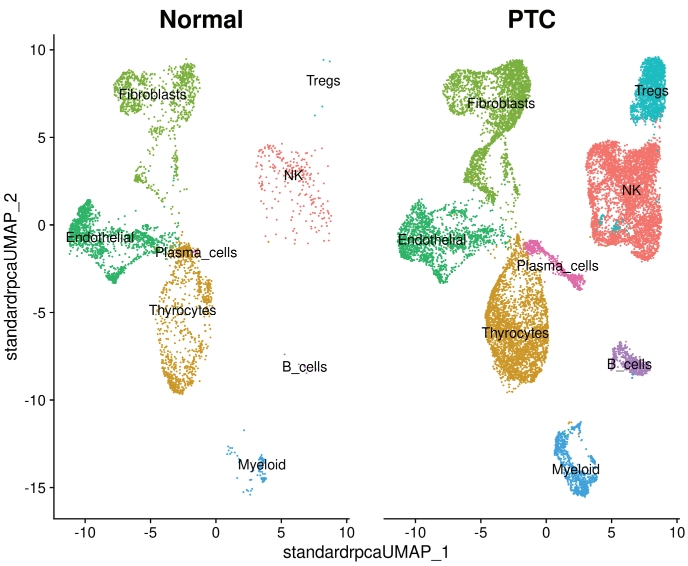

UMAP (experiment)

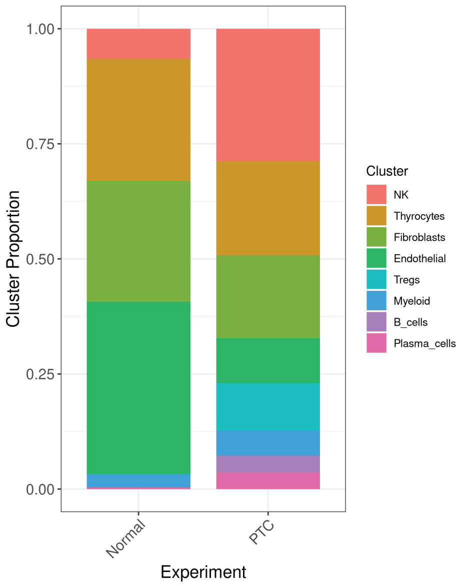

Experiment Cluster Proportions

**Show** 10 **entries** **Search:**

Table 2: Number of cells (Proportion of cells) in each Cluster by Experiment.

| | NK | Thyrocytes | Fibroblasts | Endothelial | Tregs | Myeloid | B_cells | Plasma_cells | totals |
|---|---|---|---|---|---|---|---|---|---|
| Normal | 245 (0.065) | 998 (0.266) | 983 (0.262) | 1406 (0.375) | 5 (0.001) | 100 (0.027) | 6 (0.002) | 10 (0.003) | 3753 |
| PTC | 4683 (0.288) | 3338 (0.205) | 2927 (0.18) | 1589 (0.098) | 1684 (0.103) | 881 (0.054) | 604 (0.037) | 580 (0.036) | 16286 |
| totals | 4928 | 4336 | 3910 | 2995 | 1689 | 981 | 610 | 590 | 20039 |

Showing 1 to 3 of 3 entries

Previous 1 Next

### Differentially Expressed Genes (DEGs) for each Cluster by Experiment

▼ **Table 3 Column Descriptions (CLICK TO EXPAND)**

- **gene**: gene symbol
- **cluster**: cell type cluster annotation
- **comparison**: group comparison for the analysis, formatted as a contrast where groupA is the comparison level and groupB is the reference level (groupA_vs_groupB), which indicates the directionality of the log2 fold change (avg_log2FC)
- **avg_log2FC**: log2 fold change of the average expression between the two groups being compared (see comparison column for the group comparison information)
  - *positive avg_log2FC*: gene is more highly expressed in the first group (groupA) compared to the second group (groupB)
  - *negative avg_log2FC*: gene is more highly expressed in the second group (groupB) compared to the first group (groupA)
- **pct.1**: proportion of cells expressing the gene in the first group (groupA)
- **pct.2**: proportion of cells expressing the gene in the first group (groupB)
- **p_val**: p-value from Wilcox Rank Sum test comparison of conditions
- **p_val_adj**: adjusted p-value (p_val) based on Bonferroni correction using all genes in the dataset

Column visibility | Copy | Excel | PDF | Print | **Show** 10 **entries** **Search:**

Table 3: DEGs by Cluster + Experimental Comparison.

| gene | cluster | comparison | avg_log2FC | p_val | p_val_adj | pct.1 | pct.2 |
|---|---|---|---|---|---|---|---|
| All | All | All | All | All | All | All | All |
| C16orf89 | B_cells | PTC_vs_Normal | -8.378 | 1.07e-45 | 4.09e-41 | 0 | 0.333 |
| MT1G | B_cells | PTC_vs_Normal | -8.652 | 1.07e-45 | 4.09e-41 | 0 | 0.333 |
| TFF3 | B_cells | PTC_vs_Normal | -6.665 | 3.75e-36 | 1.44e-31 | 0.018 | 0.833 |
| AC100803.3 | B_cells | PTC_vs_Normal | -7.227 | 8.57e-31 | 3.27e-26 | 0.002 | 0.333 |
| CCN1 | B_cells | PTC_vs_Normal | -5.924 | 2.01e-29 | 7.68e-25 | 0.007 | 0.5 |
| KIAA1522 | B_cells | PTC_vs_Normal | -7.859 | 1.29e-23 | 4.92e-19 | 0 | 0.167 |
| LINC01140 | B_cells | PTC_vs_Normal | -7.691 | 1.29e-23 | 4.92e-19 | 0 | 0.167 |
| GSTM1 | B_cells | PTC_vs_Normal | -7.619 | 1.29e-23 | 4.92e-19 | 0 | 0.167 |
| AC121247.1 | B_cells | PTC_vs_Normal | -8.21 | 1.29e-23 | 4.92e-19 | 0 | 0.167 |
| ANXA3 | B_cells | PTC_vs_Normal | -8.21 | 1.29e-23 | 4.92e-19 | 0 | 0.167 |

Showing 1 to 10 of 19,079 entries

Previous 1 2 3 4 5 … 1,908 Next

#### Pathway Analysis (Gene Set Enrichment Analysis) for each Cluster by Experiment

##### Ranked by adjusted.p.value

▼ **Table 4 Column Descriptions (CLICK TO EXPAND)**

- **pathway**: pathway gene set name
- **database**: the Molecular Signatures Database (MSigDB) collection of the indicated pathway (Description), where hallmark is the Hallmark collection of gene sets (H collection), curated is the Curated collection of gene sets (C2 collection), and ontology is the Ontology collection of gene sets (C5 collection)
- **cluster**: cell type cluster annotation
- **comparison**: group comparison for the analysis, formatted as a contrast where groupA is the comparison level and groupB is the reference level (groupA_vs_groupB), which indicates the directionality of the ES and NES columns
- **ES**: enrichment score, which reflects the overrpresenation of genes from the pathway gene set at the top (positive ES) or bottom (negative ES) of the list of differentially expressed genes (ranked by adjusted p-value)
  - *positive ES (and NES)*: pathway is enriched in the first group (groupA) compared to the second group (groupB) (see comparison column for information on groups)
  - *negative ES (and NES)*: pathway is enriched in the second group (groupB) compared to the first group (groupA) (see comparison column for information on groups)
- **NES**: normalized enrichment score (ES), which is the enrichment score normalized to the mean enrichment of random samples of the same size (see ES description for directionality information)
- **p-adjust**: Benjamini-Hochberg (BH) adjusted p-value from gene set enrichment analysis (GSEA) for the normalized enrichment score (NES)
- **leadingEdge**: genes in the leading edge of the analysis (genes most responsible for driving the enrichment of the pathway)
- **size**: the number of genes in the pathway gene set
- **leadingEdge_geneCount**: number of genes in the leadingEdge

Column visibility ▼ | Copy | Excel | PDF | Print | **Show** 10 **entries** **Search:**

Table 4: Pathway Analysis results from gene set enrichment analysis (GSEA) using fgsea with adjusted p-value as the ranking metric.

| pathway | database | cluster | comparison | ES | NES | p.adjust | leadingEdge | size | leadingEdge_geneCount |
|---|---|---|---|---|---|---|---|---|---|
| All | All | All | All | All | All | All | All | All | All |
| HALLMARK COMPLEMENT | hallmark | B_cells | PTC_vs_Normal | 0.314 | 2.399 | 0.021 | LCK\|CLU\|HSPA1A\|CEBPB\|S100A13\|GNB2\|JAK2\|C1S\|ZEB1\|AKAP10\|RNF4\|PLAUR\|FYN\|KLK1\|PIK3CG\|GCA\|PIK3CA\|RHOG\|L3MBTL4\|USP8\|RABIF\|PSEN1\|DOCK10\|CR2\|CBLB\|MAFF\|ITGAM\|PRCP\|TIMP2\|STX4\|SH2B3\|FN1\|LGALS3\|PIM1\|KIF2A\|DOCK9\|PLSCR1\|KYNU\|CASP7 | 47 | 39 |
| GOBP MITOCHONDRION ORGANIZATION | ontology | Endothelial | PTC_vs_Normal | 0.988 | 1.163 | 0.032 | FMC1\|DNAJC19\|TIMM22\|ACAA2\|UQCC3\|COX7A2 | 47 | 6 |
| GOCC MITOCHONDRIAL PROTEIN CONTAINING COMPLEX | ontology | Endothelial | PTC_vs_Normal | 0.992 | 1.254 | 0.097 | MRPS18A\|DNAJC19\|TIMM22\|ATP5MC3 | 26 | 4 |
| NABA CORE MATRISOME | curated | Fibroblasts | PTC_vs_Normal | 0.979 | 1.202 | 0.040 | LTBP1\|LAMA3\|POSTN\|SVEP1 | 52 | 4 |
| NABA ECM GLYCOPROTEINS | curated | Fibroblasts | PTC_vs_Normal | 0.984 | 1.283 | 0.040 | LTBP1\|LAMA3\|POSTN\|SVEP1 | 34 | 4 |
| GOMF CALCIUM ION BINDING | ontology | Fibroblasts | PTC_vs_Normal | 0.985 | 1.173 | 0.081 | DGKG\|RYR2\|LTBP1\|EGFL6\|SVEP1 | 65 | 5 |
| GOBP PEPTIDE ANTIGEN ASSEMBLY WITH MHC CLASS II PROTEIN COMPLEX | ontology | Myeloid | PTC_vs_Normal | 0.985 | 3.268 | 0.000 | HLA-DQB2\|HLA-DOA | 12 | 2 |
| GOBP PEPTIDE ANTIGEN ASSEMBLY WITH MHC PROTEIN COMPLEX | ontology | Myeloid | PTC_vs_Normal | 0.985 | 3.268 | 0.000 | HLA-DQB2\|HLA-DOA | 12 | 2 |
| GOCC MHC CLASS II PROTEIN COMPLEX | ontology | Myeloid | PTC_vs_Normal | 0.983 | 3.355 | 0.000 | HLA-DQB2\|HLA-DOA | 13 | 2 |
| GOCC MHC PROTEIN COMPLEX | ontology | Myeloid | PTC_vs_Normal | 0.983 | 3.355 | 0.000 | HLA-DQB2\|HLA-DOA | 13 | 2 |

Showing 1 to 10 of 12 entries

Previous 1 2 Next

#### Ranked by avglog2FC

▼ **Table 5 Column Descriptions (CLICK TO EXPAND)**

- **pathway**: pathway gene set name
- **database**: the Molecular Signatures Database (MSigDB) collection of the indicated pathway (Description), where hallmark is the Hallmark collection of gene sets (H collection), curated is the Curated collection of gene sets (C2 collection), and ontology is the Ontology collection of gene sets (C5 collection)
- **cluster**: cell type cluster annotation
- **comparison**: group comparison for the analysis, formatted as a contrast where groupA is the comparison level and groupB is the reference level (groupA_vs_groupB), which indicates the directionality of the ES and NES columns
- **ES**: enrichment score, which reflects the overrpresenation of genes from the pathway gene set at the top (positive ES) or bottom (negative ES) of the list of differentially expressed genes (ranked by average log2 fold change)
  - *positive ES (and NES)*: pathway is enriched in the first group (groupA) compared to the second group (groupB) (see comparison column for information on groups)
  - *negative ES (and NES)*: pathway is enriched in the second group (groupB) compared to the first group (groupA) (see comparison column for information on groups)
- **NES**: normalized enrichment score (ES), which is the enrichment score normalized to the mean enrichment of random samples of the same size (see ES description for directionality information)
- **p-adjust**: Benjamini-Hochberg (BH) adjusted p-value from gene set enrichment analysis (GSEA) for the normalized enrichment score (NES)
- **leadingEdge**: genes in the leading edge of the analysis (genes most responsible for driving the enrichment of the pathway)
- **size**: the number of genes in the pathway gene set
- **leadingEdge_geneCount**: number of genes in the leadingEdge

Column visibility ▼ | Copy | Excel | PDF | Print | **Show** 10 **entries** **Search:**

Table 5: Pathway Analysis results from gene set enrichment analysis (GSEA) using fgsea with average log2 fold change as the ranking metric.

| pathway | database | cluster | comparison | ES | NES | p.adjust | leadingEdge | size | leadingEdge_geneCount |
|---|---|---|---|---|---|---|---|---|---|
| All | All | , | All | | | A | All | | All |
| NABA CORE MATRISOME | curated | B_cells | PTC_vs_Normal | -0.479 | -1.736 | 0.029 | EFEMP1\|CCN1\|IGFBP5\|NPNT\|TINAGL1\|VWF\|SPON2\|SPON1\|FGL2\|COL9A3\|INTS6L | 23 | 11 |
| NABA ECM GLYCOPROTEINS | curated | B_cells | PTC_vs_Normal | -0.508 | -1.759 | 0.029 | EFEMP1\|CCN1\|IGFBP5\|NPNT\|TINAGL1\|VWF\|SPON2\|SPON1 | 20 | 8 |
| NABA ECM REGULATORS | curated | B_cells | PTC_vs_Normal | -0.547 | -1.776 | 0.022 | ITIH5\|BMP1\|P4HA2\|TLL2\|TIMP3\|P3H3\|SERPINF1\|TIMP2\|EGLN2 | 16 | 9 |
| NABA MATRISOME | curated | B_cells | PTC_vs_Normal | -0.445 | -2.096 | 0.000 | ANXA3\|BMP2\|FGF5\|ITIH5\|EFEMP1\|CCN1\|MUC12\|VEGFC\|IGFBP5\|BMP1\|NPNT\|TINAGL1\|P4HA2\|TLL2\|SDC2\|TIMP3\|FGF18\|P3H3\|VWF\|S100A13\|SPON2\|SERPINF1\|SPON1\|OSM\|SEMA7A\|IL10\|TIMP2\|FGL2\|EGLN2\|COL9A3 | 76 | 30 |
| NABA MATRISOME ASSOCIATED | curated | B_cells | PTC_vs_Normal | -0.445 | -1.965 | 0.003 | ANXA3\|BMP2\|FGF5\|ITIH5\|MUC12\|VEGFC\|BMP1\|P4HA2\|TLL2\|SDC2\|TIMP3\|FGF18\|P3H3\|S100A13\|SERPINF1\|OSM\|SEMA7A\|IL10\|TIMP2 | 53 | 19 |
| NABA SECRETED FACTORS | curated | B_cells | PTC_vs_Normal | -0.438 | -1.586 | 0.062 | BMP2\|FGF5\|VEGFC\|FGF18\|S100A13\|OSM\|IL10\|S100A4\|CLCF1\|IL23A\|TNFSF10\|HCFC2\|S100A6\|EBI3 | 23 | 14 |
| SIG BCR SIGNALING PATHWAY | curated | B_cells | PTC_vs_Normal | 0.543 | 2.548 | 0.000 | PPP3CC\|INPP5D\|AKT3\|GSK3B\|SOS2\|PPP1R13B\|BLNK\|PLCG2\|PPP3CB\|CR2\|NFATC2\|PPP3CA\|CSK\|CD19\|MAP4K1\|NFATC1\|BTK | 21 | 17 |
| SIG PIP3 SIGNALING IN B LYMPHOCYTES | curated | B_cells | PTC_vs_Normal | 0.472 | 1.853 | 0.022 | AKT3\|RPS6KA3\|PPP1R13B\|PLCG2\|PTEN\|GAB1\|TEC\|CD19\|BTK | 13 | 9 |
| HALLMARK COAGULATION | hallmark | B_cells | PTC_vs_Normal | -0.521 | -1.847 | 0.099 | GNG12\|BMP1\|RGN\|TIMP3\|VWF\|S100A13\|CFH\|RABIF | 21 | 8 |
| HALLMARK MITOTIC SPINDLE | hallmark | B_cells | PTC_vs_Normal | 0.364 | 2.62 | 0.000 | SSH2\|RAPGEF6\|RAPGEF5\|DOCK2\|RAB3GAP1\|MYO9B\|NCK2\|ABR\|DLG1\|SPTAN1\|RICTOR\|TRIO\|RABGAP1\|NEDD9\|NF1\|RANBP9\|FGD6\|MYO1E\|RASA1\|ABI1\|EPB41L2\|CLASP1\|LRPPRC\|NOTCH2\|SMC4\|CLIP2 | 69 | 26 |

Showing 1 to 10 of 853 entries

Previous 1 2 3 4 5 … 86 Next

### Z-Scores

Z-score transformation of gene expression data standardizes the values by converting them to a common scale with a mean of 0 and a standard deviation of 1.

- **Positive z-score**: gene with higher gene expression in the specific experiment/sample comapred to the mean
- **Negative z-scores**: gene with lower gene expression in the specific experiment/sample comapred to the mean

Z-scores have been calculated for the following groupings:

- Cluster (Table 6)
- Cluster and Experiment (Table 7)
- Cluster, Sample, and Experiment (Table 8)

Z-scores allow for comparison of gene expression by removing biases due to differences in scale or distribution. This makes it easier to identify relative changes in expression while also granting the ability to see which genes are most highly expressed within each column in the tables.

##### Split by Cluster

Column visibility | Copy | Excel | PDF | Print **Show** 10 **entries** **Search:**

Table 6: Z-Scores by Cluster.

| gene | NK | Thyrocytes | Fibroblasts | Endothelial | Tregs | Myeloid | B.cells | Plasma.cells |
|---|---|---|---|---|---|---|---|---|
| All | All | All | All | All | All | All | All | All |
| MIR1302-2HG | -0.083 | -0.067 | -0.069 | -0.083 | -0.087 | -0.099 | -0.086 | -0.013 |
| FAM138A | -0.083 | -0.067 | -0.069 | -0.083 | -0.087 | -0.099 | -0.086 | -0.013 |
| OR4F5 | -0.083 | -0.067 | -0.069 | -0.083 | -0.087 | -0.099 | -0.086 | -0.013 |
| AL627309.1 | -0.083 | -0.066 | -0.068 | -0.081 | -0.083 | -0.096 | -0.083 | -0.013 |
| AL627309.3 | -0.083 | -0.067 | -0.069 | -0.083 | -0.087 | -0.099 | -0.086 | -0.013 |
| AL627309.2 | -0.083 | -0.067 | -0.069 | -0.083 | -0.087 | -0.099 | -0.086 | -0.013 |
| AL627309.4 | -0.083 | -0.067 | -0.069 | -0.083 | -0.087 | -0.099 | -0.086 | -0.013 |
| AL732372.1 | -0.083 | -0.066 | -0.069 | -0.083 | -0.087 | -0.099 | -0.086 | -0.013 |
| OR4F29 | -0.083 | -0.067 | -0.069 | -0.083 | -0.087 | -0.099 | -0.086 | -0.013 |
| AC114498.1 | -0.083 | -0.067 | -0.069 | -0.083 | -0.087 | -0.099 | -0.086 | -0.013 |

Showing 1 to 10 of 38,224 entries Previous 1 2 3 4 5 … 3,823 Next

##### Split by Cluster + Experiment

Column visibility | Copy | Excel | PDF | Print **Show** 10 **entries** **Search:**

Table 7: Z-Scores by Cluster + Exerimental Condition.

| gene | NK_Normal | NK_PTC | Thyrocytes_Normal | Thyrocytes_PTC | Fibroblasts_Normal | Fibroblasts_PTC | Endothelial_Normal | Endothelial_PTC | Tregs_Normal | Tregs_PTC | Myeloid_Normal | Myeloid_PTC | B.cells_Normal | B.cells_PTC | Plasma.cells_Normal | Plasma.cells_PTC |
|---|---|---|---|---|---|---|---|---|---|---|---|---|---|---|---|---|
| All | All | All | All | All | All | All | All | All | All | All | All | All | All | All | All | All |
| MIR1302-2HG | -0.06 | -0.085 | -0.051 | -0.072 | -0.067 | -0.069 | -0.084 | -0.082 | -0.072 | -0.087 | -0.088 | -0.1 | -0.07 | -0.086 | -0.009 | -0.013 |
| FAM138A | -0.06 | -0.085 | -0.051 | -0.072 | -0.067 | -0.069 | -0.084 | -0.082 | -0.072 | -0.087 | -0.088 | -0.1 | -0.07 | -0.086 | -0.009 | -0.013 |
| OR4F5 | -0.06 | -0.085 | -0.051 | -0.071 | -0.067 | -0.069 | -0.084 | -0.082 | -0.072 | -0.087 | -0.088 | -0.1 | -0.07 | -0.086 | -0.009 | -0.013 |
| AL627309.1 | -0.06 | -0.084 | -0.051 | -0.071 | -0.067 | -0.068 | -0.082 | -0.08 | -0.072 | -0.084 | -0.088 | -0.096 | -0.07 | -0.083 | -0.009 | -0.013 |
| AL627309.3 | -0.06 | -0.085 | -0.051 | -0.072 | -0.067 | -0.069 | -0.084 | -0.081 | -0.072 | -0.087 | -0.088 | -0.1 | -0.07 | -0.086 | -0.009 | -0.013 |
| AL627309.2 | -0.06 | -0.085 | -0.051 | -0.072 | -0.067 | -0.069 | -0.084 | -0.082 | -0.072 | -0.087 | -0.088 | -0.1 | -0.07 | -0.086 | -0.009 | -0.013 |
| AL627309.4 | -0.06 | -0.085 | -0.051 | -0.072 | -0.067 | -0.069 | -0.084 | -0.082 | -0.072 | -0.087 | -0.088 | -0.1 | -0.07 | -0.086 | -0.009 | -0.013 |
| AL732372.1 | -0.06 | -0.085 | -0.051 | -0.071 | -0.067 | -0.069 | -0.084 | -0.082 | -0.072 | -0.087 | -0.088 | -0.1 | -0.07 | -0.086 | -0.009 | -0.013 |
| OR4F29 | -0.06 | -0.085 | -0.051 | -0.072 | -0.067 | -0.069 | -0.084 | -0.082 | -0.072 | -0.087 | -0.088 | -0.1 | -0.07 | -0.086 | -0.009 | -0.013 |
| AC114498.1 | -0.06 | -0.085 | -0.051 | -0.071 | -0.067 | -0.069 | -0.084 | -0.082 | -0.072 | -0.087 | -0.088 | -0.1 | -0.07 | -0.086 | -0.009 | -0.013 |

Showing 1 to 10 of 38,224 entries Previous 1 2 3 4 5 … 3,823 Next

##### Split by Cluster + Experiment + Sample

Column visibility | Copy | Excel | PDF | Print **Show** 10 **entries** **Search:**

Table 7: Z-Scores by Cluster + Exerimental Condition.

| gene | NK_Normal_NT | NK_PTC_T1L | NK_PTC_T1R | NK_PTC_T2L | NK_PTC_T2R | NK_PTC_T3L | NK_PTC_T3R | Thyrocytes_Normal_NT | Thyrocytes_PTC_T1L | Thyrocytes_PTC_T1R | Thyrocytes_PTC_T2L | Thyrocytes_PTC_T2R | Thyrocytes_PTC_T3L | Thyrocytes_PTC_T3R | Fibroblasts_Normal_NT |
|---|---|---|---|---|---|---|---|---|---|---|---|---|---|---|---|
| All | All | All | All | All | All | All | All | All | All | All | All | All | All | All | All |
| MIR1302-2HG | -0.06 | -0.051 | -0.046 | -0.106 | -0.094 | -0.1 | -0.095 | -0.051 | -0.031 | -0.055 | -0.071 | -0.104 | -0.082 | -0.071 | -0.067 |
| FAM138A | -0.06 | -0.051 | -0.046 | -0.106 | -0.094 | -0.1 | -0.095 | -0.051 | -0.031 | -0.055 | -0.071 | -0.104 | -0.082 | -0.071 | -0.067 |
| OR4F5 | -0.06 | -0.051 | -0.046 | -0.106 | -0.094 | -0.1 | -0.095 | -0.051 | -0.031 | -0.048 | -0.071 | -0.104 | -0.082 | -0.071 | -0.067 |
| AL627309.1 | -0.06 | -0.051 | -0.046 | -0.105 | -0.093 | -0.1 | -0.095 | -0.051 | -0.031 | -0.055 | -0.063 | -0.104 | -0.082 | -0.07 | -0.067 |
| AL627309.3 | -0.06 | -0.051 | -0.046 | -0.106 | -0.094 | -0.1 | -0.095 | -0.051 | -0.031 | -0.055 | -0.071 | -0.104 | -0.082 | -0.071 | -0.067 |
| AL627309.2 | -0.06 | -0.051 | -0.046 | -0.106 | -0.094 | -0.1 | -0.095 | -0.051 | -0.031 | -0.055 | -0.071 | -0.104 | -0.082 | -0.071 | -0.067 |
| AL627309.4 | -0.06 | -0.051 | -0.046 | -0.106 | -0.094 | -0.1 | -0.095 | -0.051 | -0.031 | -0.055 | -0.071 | -0.104 | -0.082 | -0.071 | -0.067 |
| AL732372.1 | -0.06 | -0.051 | -0.046 | -0.106 | -0.094 | -0.1 | -0.095 | -0.051 | -0.031 | -0.055 | -0.071 | -0.104 | -0.081 | -0.07 | -0.067 |
| OR4F29 | -0.06 | -0.051 | -0.046 | -0.106 | -0.094 | -0.1 | -0.095 | -0.051 | -0.031 | -0.055 | -0.071 | -0.104 | -0.082 | -0.071 | -0.067 |
| AC114498.1 | -0.06 | -0.051 | -0.046 | -0.106 | -0.094 | -0.1 | -0.095 | -0.051 | -0.031 | -0.055 | -0.071 | -0.104 | -0.082 | -0.071 | -0.067 |

Showing 1 to 10 of 38,224 entries Previous 1 2 3 4 5 … 3,823 Next

[Scrollable table in HTML report]

#### UMAP by Cell Cycling Phase

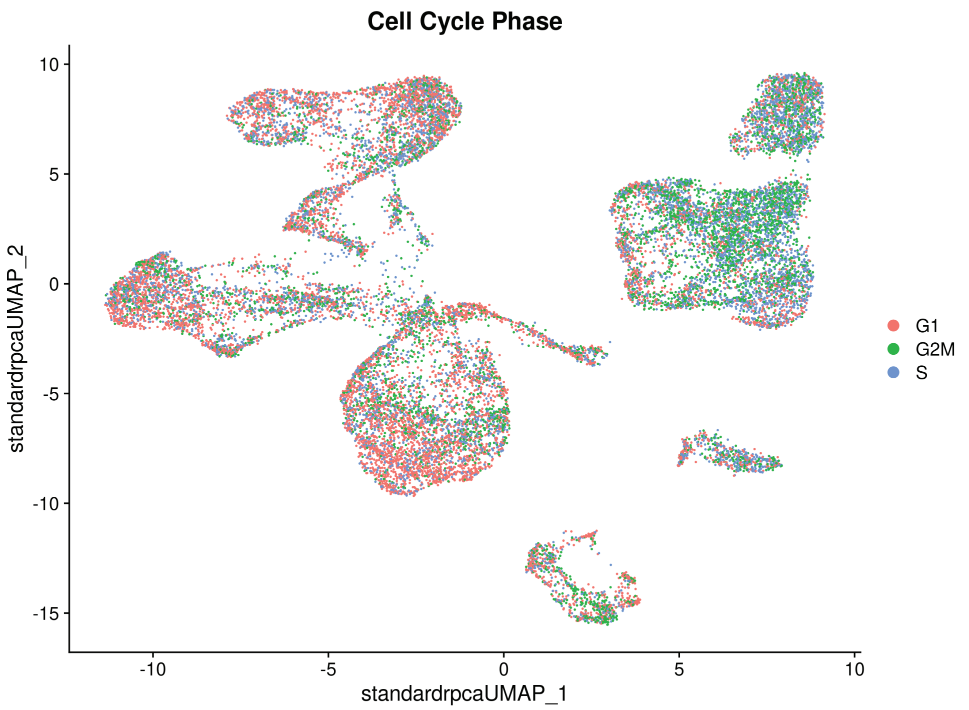

UMAP (Cell Cycle Phase)

#### Variable Genes

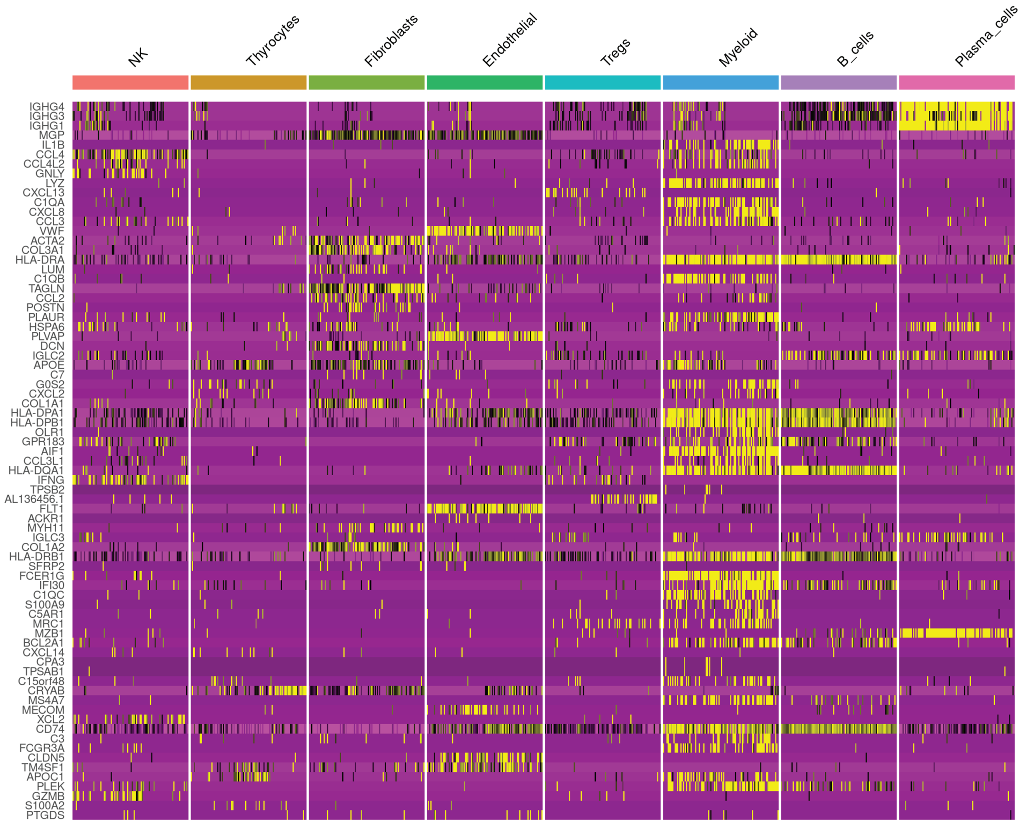

Copy Excel Print

**Show** 10 **entries**

**Search:**

Table 9: Variable Genes.

| **genes** |
|---|
| IGHG4 |
| IGHG3 |
| IGHG1 |
| MGP |
| IL1B |
| CCL4 |
| CCL4L2 |
| GNLY |
| LYZ |
| CXCL13 |

Showing 1 to 10 of 3,000 entries

Previous 1 2 3 4 5 … 300 Next

Heatmap Plot of Variable Genes

### Conserved Genes

**For all experimental groups, the combined miminum p-value = 0.**
Conserved genes are genes tha are conserved between groups (Experiment) within a cluster.
NOTE: If there is single body split into multiple clusters, conserved markers for those clusters are UNlikely to be genes used for annotation purposes.

Column visibility ▼ Copy Excel PDF Print **Show** 10 **entries** **Search:**

Table 10: Conserved Genes by Across Exerimental Conditions by Cluster.

| gene | cluster |
|---|---|
| All | All |
| CD79A | B_cells |
| IGKC | B_cells |
| MS4A1 | B_cells |
| SPIB | B_cells |
| JCHAIN | B_cells |
| CXCR5 | B_cells |
| IGHA1 | B_cells |
| NIBAN3 | B_cells |
| LINC01781 | B_cells |
| FCRL2 | B_cells |

Showing 1 to 10 of 2,283 entries Previous 1 2 3 4 5 … 229 Next

Conserved_Genes

### Trajectory Analysis (Monocle3)

**Please see the docuemntation for Monocle3 (https://cole-trapnell-lab.github.io/monocle3/) for additional information and instructions on trajectory analysis and how to read the trajectory graph.**
In Monocle3, the trajectory graph edges and nodes are visualized on top of a UMAP. There are two different types of nodes that are present on the graph, here visualized as:

- **Light gray circles**: "leaf" nodes, which correspond to a different cell fate outcome of the trajectory
- **Black circles**: "branch" nodes, which are nodes from which cells may travel to different possible connected leaf nodes.

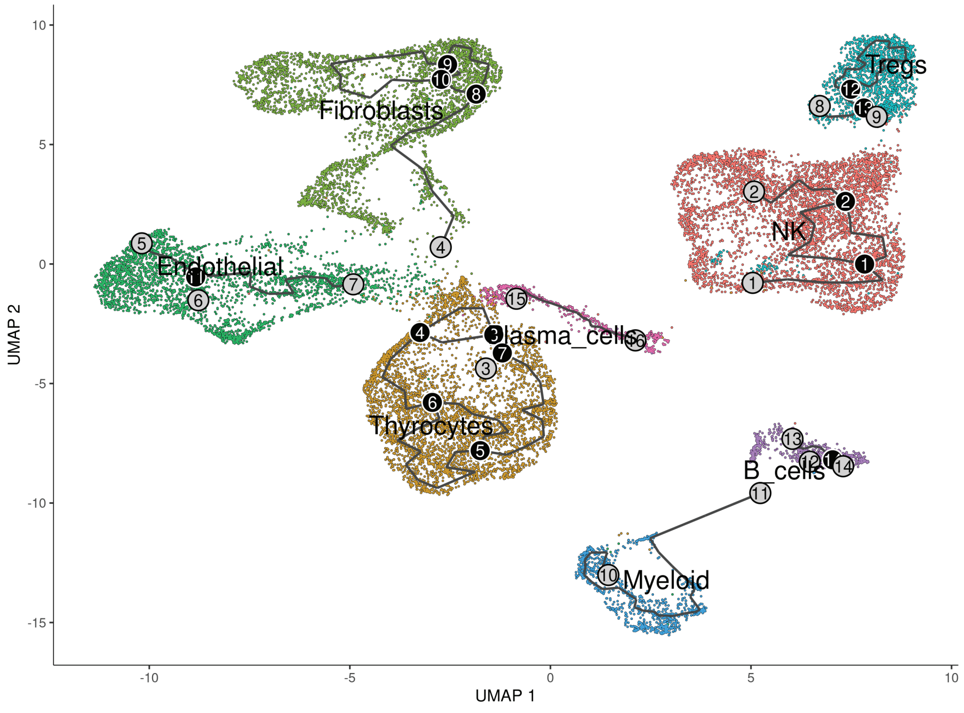

Trajectory Analysis

### 4 Supplementary PDF 4: Benchmarking Report

### prjna790856

#### Benchmarking Report

2025-03-23 18:10:54

### Pipeline Information

SWANS, version 2.0

### YAML Configuration Parameters

##### Species

Organism: human

##### Initial Processing

Was CellRanger Run? FALSE
If yes, were bam files created? FALSE
Was a MultiQC report created with CellRanger output? FALSE
Was SoupX Run? FALSE
Was DoubletFinder Run? TRUE

How many PCA components were initially investigated? 30
How may variable features were used? 3000
How was the data scaled? variable

Were mitochondira genes regressed out? FALSE
Were ribosomal genes regressed out? FALSE
Were cell cycling genes regressed out? FALSE
Name of user-supplied file containing additional genes to regress out?

Normalization Method(s): sct,standard
Integration Method(s): cca,harmony,rpca
Resolution(s): 0.1,0.2,0.3

Reference-Based Integration? FALSE
Reference-Based Samples:

Was Azimuth Run? FALSE
Were lables transferred from an additional seurat object? FALSE
Was Azimuth Run? FALSE

Was FindConservedMarkers run? FALSE
Was tSNE used? FALSE

How was the data stored?
How many threads were used for parallel processing? 30
Max assigned memory? NA

Did user supply additional genes for visualization? gene_files/prjna1185392.txt
How did they want the additional genes visualized? dot

---

##### Final Processing

Was Trajectory Analysis Run? TRUE
How many threads were used for parallel processing? 30
How was the final data stored? cellchat
Were conserved genes found? TRUE
Name of user-supplied file containing (final) genes for visualization?
How were they visualized?

### Sample Information

Show 20 entries

Search:

Table 1: Sample Data.

| | samples | condition |
|---|---|---|
| 1 | T1L | PTC |
| 2 | T1R | PTC |
| 3 | T2L | PTC |
| 4 | T2R | PTC |
| 5 | T3L | PTC |
| 6 | T3R | PTC |
| 7 | NT | Normal |

Showing 1 to 7 of 7 entries

Previous 1 Next

### Benchmarks

Column visibility Copy Excel PDF Print Show 30 entries

Search:

Table 2: Benchmarking results for all Snakemake Rule

| h.m.s | max_rss | max_vms | io_in | io_out | mean_load | cpu_time | benchmark_application |
|---|---|---|---|---|---|---|---|
| | All | All | | | All | All | All |
| 0:00:02 | 294.79 | 1049461.6 | 0 | 0.03 | 51.66 | 1.9 | prjna790856_benchmark_report |
| 0:00:02 | 204.55 | 745.8 | 0.06 | 0.55 | 66.35 | 2.33 | prjna790856_benchmark_table |
| 0:27:20 | 10768.06 | 11508.39 | 1382.87 | 547.49 | 76.63 | 1257.34 | prjna790856_cluster_plots |
| 0:01:38 | 3054.05 | 3661.14 | 0.04 | 10.43 | 75.2 | 74.7 | prjna790856_create_initial_seurat |
| 0:09:20 | 302612.18 | 339064.77 | 18.7 | 1646.1 | 94.39 | 543.13 | prjna790856_final_analysis |
| 0:00:31 | 1137.22 | 1049773.05 | 41.29 | 32.32 | 46.58 | 15.66 | prjna790856_final_report |
| 0:00:00 | 2.82 | 12.64 | 0 | 0 | 0 | 0 | prjna790856_memory |
| 0:23:19 | 14354.44 | 16025.03 | 0 | 5.8 | 94.1 | 1317.92 | prjna790856_NT_doubletFinder |
| 0:00:04 | 323.53 | 1049446.15 | 4.33 | 18.45 | 33.47 | 2.29 | prjna790856_qc_report |
| 1:13:52 | 11649.82 | 30808.05 | 66.29 | 714.7 | 156.32 | 6929.67 | prjna790856_seurat_analysis |
| 0:17:02 | 10261.71 | 12017.73 | 0 | 4.75 | 86.04 | 881.9 | prjna790856_T1L_doubletFinder |
| 0:13:05 | 8040.65 | 9703.91 | 0.02 | 4.71 | 73.14 | 575.9 | prjna790856_T1R_doubletFinder |

| h.m.s | max_rss | max_vms | io_in | io_out | mean_load | cpu_time | benchmark_application |
|---|---|---|---|---|---|---|---|
| 0:19:21 | 10984.89 | 12656.3 | 0 | 5.26 | 69.95 | 812.64 | prjna790856_T2L_doubletFinder |
| 0:17:21 | 9182.51 | 11105.23 | 0 | 4.59 | 76.75 | 800.63 | prjna790856_T2R_doubletFinder |
| 0:27:38 | 13498.6 | 15169.81 | 0 | 5.55 | 81.52 | 1353.05 | prjna790856_T3L_doubletFinder |
| 0:34:18 | 17726 | 19393.21 | 0 | 6.73 | 82.47 | 1698.4 | prjna790856_T3R_doubletFinder |
| 0:01:28 | 5694.97 | 6440.67 | 1314.63 | 0 | 79.97 | 71.29 | prjna790856_trajectory_analysis |

Showing 1 to 17 of 17 entries

Previous 1 Next

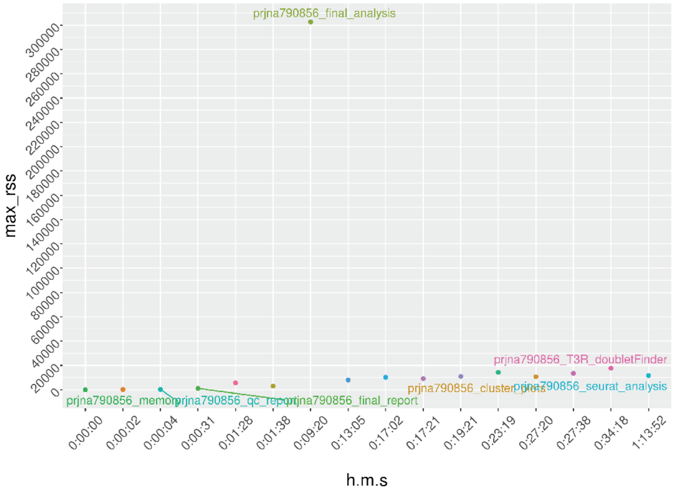

Figure 1: Running time (hour:minutes:seconds) vs Max Physical Memory

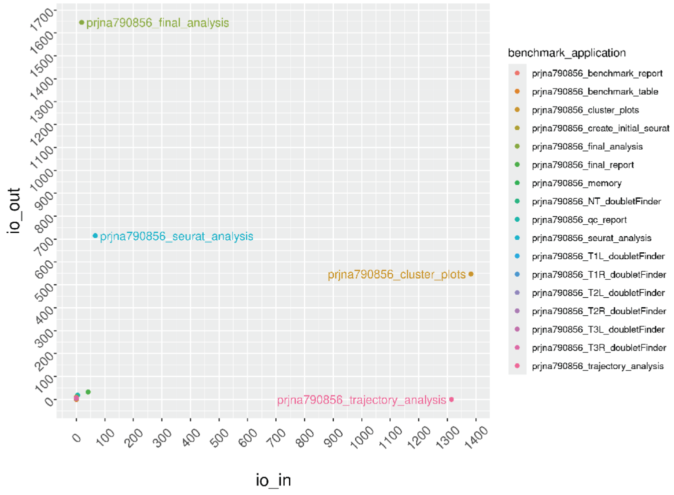

Figure 2: Number of MB read vs Number of MB written

#### psutil description

| colname | type (unit) | description |
|---|---|---|
| s | float (seconds) | Running time in seconds |
| h:m:s | string (-) | Running time in hour, minutes, seconds format |
| max_rss | float (MB) | Maximum “Resident Set Size”, this is the non-swapped physical memory a process has used. |
| max_vms | float (MB) | Maximum “Virtual Memory Size”, this is the total amount of virtual memory used by the process |
| max_uss | float (MB) | “Unique Set Size”, this is the memory which is unique to a process and which would be freed if the process was terminated right now. |
| max_pss | float (MB) | “Proportional Set Size”, is the amount of memory shared with other processes, accounted in a way that the amount is divided evenly between the processes that share it (Linux only) |
| io_in | float (MB) | the number of MB read (cumulative). |
| io_out | float (MB) | the number of MB written (cumulative). |

| colname | type (unit) | description |
|---|---|---|
| mean_load | float (-) | CPU usage over time, divided by the total running time (first row) |
| cpu_time | float(-) | CPU time summed for user and system |
